## Supplementary Notes, Figures and Tables for "A digital twin for DNA data storage based on comprehensive quantification of errors and biases"

### Supplementary Note 1

##### Details on the error analysis from sequencing reads

To estimate error rates from sequencing reads, we first mapped every sequencing read to its respective reference sequence, and then derived error rates as a function of position, base, read, and error length from all mappings. For all analyses, at most 1 million reads (including both forward and reverse reads of a sequence, equivalent to a mean coverage of about 80 reads per sequence) were randomly selected from the sequencing datasets as a representative sample to limit computing time.

Individual reads were trimmed and mapped to their reference using the sequence alignment functions implemented in the Python (v3.10) libraries RapidFuzz (v2.6) and edlib (v1.2).[1,2] For mapping, the similarity metric implemented in RapidFuzz, defined as

$$\text{Similarity} = 1 - \frac{\text{Hamming distance between read and reference}}{\text{Length of reference} + \text{Length of read}}$$

was used to select the highest-scoring reference as the appropriate reference sequence for every read. Using the metric of similarity, sequencing reads were also categorized as “matches”, “partial matches” and “non-matches”. Appropriate values for these similarity cut-offs were chosen based on the observed distribution of similarities for an experimental sequencing run compared to that of randomly generated sequences (see Supplementary Figure 1). As a result, sequences above 85% similarity were considered as matches, and sequences below 70% similarity as non-matches.

To derive error rates from the mapped sequencing reads, point mutations were categorized as substitutions, insertions, or deletions and information about each mutations’ position, involved bases, and read type were collected. To remove artefacts from mapping, some mutations were removed:

- excess deletions from the end of the mapping, present when the reference sequence is longer than the read length of the sequencer,
- misassigned deletions at the start of the mapping, present when the first base of the read is a substitution but is instead considered as a deletion,
- excess insertions from the start or the end of the mapping, which might occur when deletions decrease the total length of the read sequence.

Across all analysed sequences, error rates were then estimated from the number of mutations relative to the total number of bases in the sample. Error rates were stratified by match category (i.e., matches, partial matches, or non-matches), position within the read, position within the reference sequence, involved base, and length of the mutation. For further analysis of error rates, only those rates estimated from reads categorized as matches (similarity >85%) were used.

The complete source code and examples are publicly available in the GitHub repository.

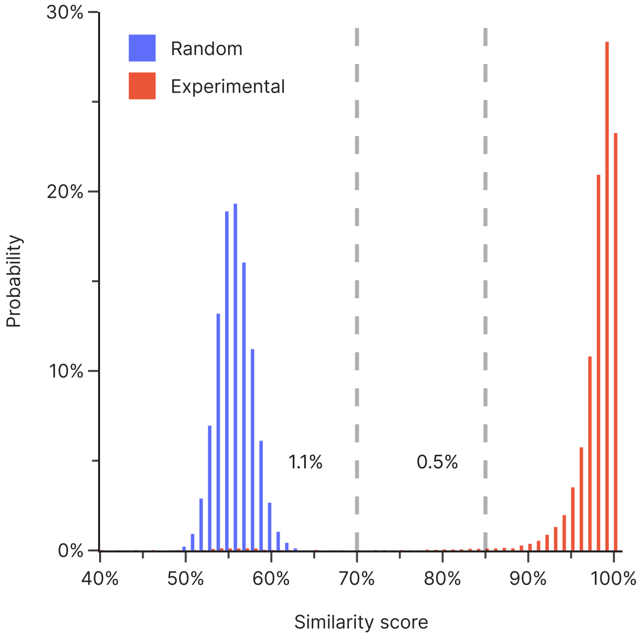


**Supplementary Figure 1: Definition of the similarity cut-off.** A similarity cut-off of 85% was chosen for the analysis based on the distribution of similarity scores from an experimental sequencing run (red) and an equivalent, randomly generated dataset (blue). Around 0.5% of the real sequencing reads scored between 70% and 85% similarity, and only 1.1% scored below 70%. In contrast, virtually all randomly generated sequences scored less than 70% similarity.

##### Details on the coverage analysis from sequencing reads

Coverage was calculated from sequencing datasets with BBMap (v38.99)[3] using the `scafstats` option for BBMap after adapter trimming with BBDuk. The complete command used was:

```
bbduk.sh -Xmx6g in1=<R1.fq.gz> in2=<R2.fq.gz> out=stdout.fq
ref=adapters ktrim=r k=23 mink=11 hdist=1 tpe tbo | bbmap.sh -Xmx6g
in=stdin.fq ref=<design_files.fasta> ordered interleaved nodisk
scafstats=scafstats.txt
```

From the generated file, the “assignedReads” column was used as abundance to calculate the normalized abundance according to:

$$\text{Normalized abundance of sequence } i = \frac{\text{Abundance of sequence } i}{\text{Mean abundance of all sequences}}$$

For determining the coverage of error-free reads only, a procedure similar to Koch et al.[4] and Erlich et al.[5] was used, consisting of read stitching using PEAR[6], filtering to remove reads with incorrect length, followed by coverage estimation using BBMap with its `perfectmode` and `scafstats` options (see also above). The commands used were:

```
pear -j 8 -f <R1.fq.gz> -r <R2.fq.gz> -o pear.all

awk '(NR%4==2 && length($0)=='$2'){print $0}'
pear.all.assembled.fastq > pear.all.good

python3 peartofasta.py <dir> # reformatting from pear output to fasta

bbmap.sh -Xmx6g in=pear.all.good.fasta ref=<design_files.fasta>
nodisk perfectmode scafstats=scafstats_pear_perfect.txt
```

As above, the “assignedReads” column from the generated file was used as abundance for the coverage analysis.

### Supplementary Note 2

#### Overview for the model of the DNA data storage process

The process model consists of three major elements: a hash map representing a pool of oligonucleotides, error generators introducing mutations into oligonucleotide pools, and classes encapsulating the error generators into the individual process steps (e.g. synthesis, PCR, …), see Supplementary Figure 2. The source code is available in the GitHub repository (see Data Availability statement).

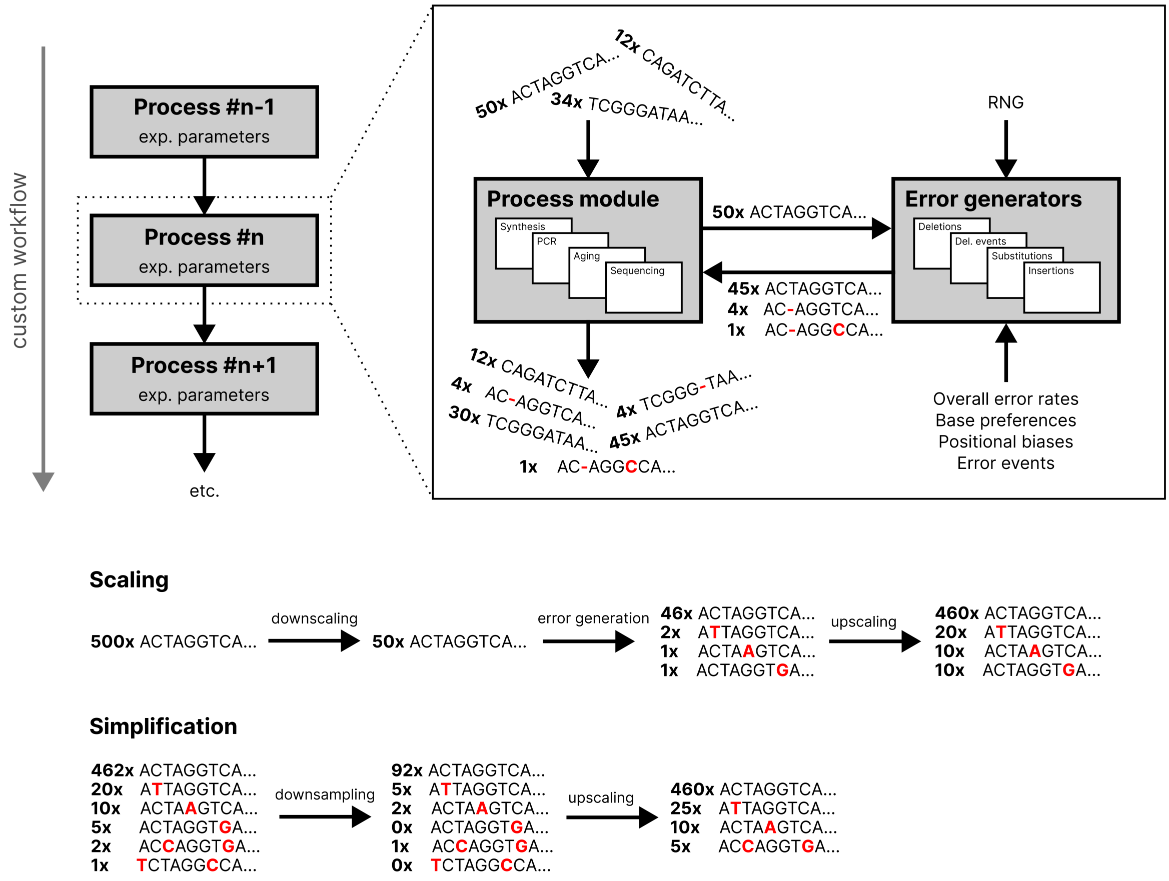


**Supplementary Figure 2: Graphical overview of the model for the DNA data storage process.** A sequence of individual process modules constitutes the user-defined, custom data storage workflow. Each process module generates erroneous sequences from its input sequences and the user-defined parameters, which are passed as input to the next process module in the workflow. Two approaches for reducing the diversity of the simulated oligo pool are used throughout the workflow. Scaling limits the number of unique sequences being generated in each process module by downscaling the oligo pools prior to error generation. Simplification also reduces pool diversity by randomly downsampling the oligo pool to generate a representative sample, before re-scaling the abundances to match the initial pool. The parameters of both approaches is selected such that the resulting pool size and diversity is orders of magnitudes larger than the sequencing read count (e.g. scaling such that around 10 new sequences are generated for each input sequence, and $10^7$ sequences as upper limit for simplification).

An oligonucleotide pool is implemented as a simple Python `dict`, mapping an oligonucleotide sequence to a value representing the number of oligos with this sequence. Basic operations, such as dilution, operate directly on the dictionary, e.g., by reducing the oligo count or removing oligo sequences. To limit the memory requirements of oligo pools, the diversity of a pool is limited by a maximum number of oligos (user-defined, $10^7$ in this study). If the total number of oligos exceeds this limit, a number of oligos equal to the limit are randomly sampled from the pool, and their abundance is upscaled to the pool's initial oligo count, …), see Supplementary Figure 2. While this effectively limits the maximum number of unique sequences ($10^7$ in this study), this limit is around 10x the usual read count, and multiple orders of magnitude larger than the number of unique sequences to be expected in our experiments (e.g., starting from only around 12000 reference sequences).

Error generators use a mean error rate, a base bias matrix, and position- and length-dependent correction factors to stochastically change the sequences from all sequences in oligo pools. In this way, substitutions, insertions, deletions, and deletion events (e.g., runs of deletions) can be introduced. In all cases, the error parameters are derived from user-supplied experimental parameters and the error analysis presented in this study. For example, the base bias matrix, representing the likelihood of a generated substitution to be of a certain type, is dependent on the type of polymerase used. Unless a non-ideal distribution of errors per read is used (e.g., for deletion events), the number of errors per oligo is given by:

$$\text{Errors in oligo} = \text{Binom}(\text{Number of nucleotides, mean error rate per nucleotide})$$

where the mean error rate per nucleotide is calculated from the position-dependent error rate and the base composition of its sequence. The position of each error is then randomly selected, with weights according to the position-dependent error rate and base preference. For substitutions and insertions, the new base is also randomly selected, with weights according to the base bias matrix. To limit the number of unique sequences being generated, the number of new erroneous sequences derived from a given input sequence is restricted…), see Supplementary Figure 2. This is achieved by scaling the oligo count for error generation to yield a specific number of new erroneous sequences on average (user-defined, 10 in this study), while enforcing an upper limit on this scaled oligo count (user-defined, $10^4$ in this study) such that small error rates are not overrepresented. After error generation, the oligo counts are rescaled to match initial counts. Based on this approach, error rates per mutation type and process step as low as $10^{-4}$ per read, or about $10^{-6}$ per nucleotide can be reasonably represented. This is more than sufficient for the error rates expected for DNA data storage applications.

Classes representing the individual process steps (e.g., synthesis, PCR, …) of the DNA data storage workflow utilize the aforementioned error generators, experimental parameters supplied by the user, and a set of process-specific settings (e.g., mean deletion rates for different synthesis providers) to model the process' impact on an oligonucleotide pool. The implementation of all process steps is described in the following subsections.

**Implementation of synthesis**

A set of reference sequences supplied by the user is used to generate an oligonucleotide pool at the specified synthesis scale. To introduce the synthesis-derived coverage bias, each sequence's oligo count is randomly chosen from a specified distribution. Following this, deletions are added to the reference sequences based on the position-, process-, and length-dependent deletion rates described in the main text. Distributional non-idealities are accounted for in the error generation by overriding the binomial error distribution (see above) with the error distribution observed experimentally. Primer regions flanking the data-encoding sequence regions are added, if desired, after error generation. This prevents the generation of oligos with errors in the primer regions, which are not amplified during PCR.

**Implementation of PCR**

To perform amplification, sequences from an oligo pool are categorized based on the presence of priming regions complementary to the specified primer sequence. The amplification of those sequences containing priming regions is modelled by a binomial process for each sequence and each cycle of amplification:

$$\text{Oligo count in cycle } i+1 = \text{ Oligo count in cycle } i + \text{Binom}(\text{Oligo count in cycle } i,\ \text{ efficiency})$$

With this binomial process, stochastic effects observed for amplification at low oligo counts[7] are taken into account. Due to the distribution of amplification efficiencies leading to PCR-induced changes in the coverage bias (see main text), each sequence has an associated, randomly chosen efficiency which is constant over a simulation run. This efficiency is also copied to all mutated oligos derived from this sequence. After amplification, substitutions are introduced at a mean rate according to:

$$\text{Mean substitution rate} = \frac{\text{Taq-polymerase error rate per cycle}}{\text{Polymerase fidelity}} * \text{Number of cycles}$$

where the polymerase fidelity is defined as the polymerase's error rate relative to Taq polymerase.[8,9] In this study, due to the use of a Taq-derived polymerase, polymerase fidelity is 1. No position- or length-related deviations for the generation of subsitutions were considered. To limit the diversity of generated sequences, only those sequences present at least 0.1% as abundant as the most common sequence, but at least 95% of all oligos, were included in the error generation.

**Implementation of storage**

The storage-induced decay of oligos is implemented as simple stochastic sampling, in-line with the findings from the main text. After sampling, substitutions are introduced as a function of the number of half-lives of decay, calculated from the fraction of intact oligos:

$$\text{Mean substitution rate} = \text{Substitution rate per half-life} * \log_2\left(\frac{1}{\text{Fraction of intact oligos}}\right)$$

No position- or length-related deviations for the generation of substitutions were considered, due to the observed insignificance of these types of errors during storage (see main text).

**Implementation of sequencing**

To perform sequencing, sequences from an oligo pool are categorized based on the presence of priming regions complementary to the specified sequencing primers. Then, oligos equal to the user-defined number of reads are randomly sampled from the pool, and substitutions are introduced based on read- and position-dependent error rates, as well as a base bias matrix. After the introduction of errors, forward and reverse reads were generated via the sequence and its complement starting from each sequencing primer region respectively. If the sequence length is smaller than the sequencing read length, random nucleotides were added to the read instead. One output file per read direction is then generated in the FASTQ format, including a quality string consisting only of “F”, the highest quality score on the Illumina iSeq 100 sequencing system.

#### Supplementary Note 3

##### Estimation of substitution errors during synthesis

Due to the experimental design of our study, the rate of substitutions during synthesis is not directly available from the sequencing data. However, the amplification experiments allow for an indirect estimation of the rate of substitutions introduced during synthesis by extrapolating the substitution rate to zero PCR cycles (see Supplementary Figure 3). This estimation yields a baseline substitution rate of $2.72 \pm 0.42 \cdot 10^{-3}$ $nt^{-1}$ (electrochemical synthesis) and $2.48 \pm 0.13 \cdot 10^{-3}$ $nt^{-1}$ (material deposition), respectively. Considering a mean substitution rate of around $1.8 \cdot 10^{-3}$ $nt^{-1}$ caused by sequencing (as estimated from PhiX-derived reads, see main text), a substitution rate of around $0.8 \cdot 10^{-3}$ $nt^{-1}$ cannot be accounted for by PCR and sequencing alone. It is likely that these substitutions arise during synthesis, potentially because of non-quantitative capping. However, in the context of DNA data storage, the contribution of these synthesis-derived substitutions to the overall error rate is minor, as they account for only around 10% of the mean number of substitutions observed across all experimental conditions.

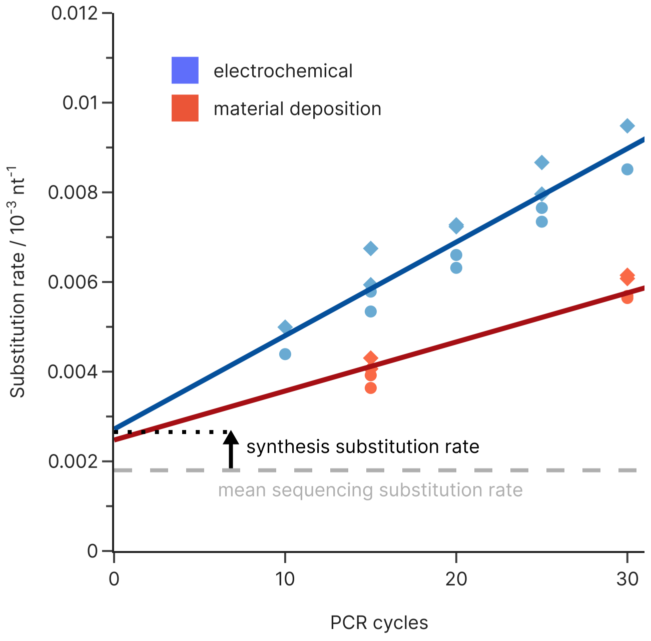


**Supplementary Figure 3: Estimation of the synthesis-derived substitution rate from amplification experiments.** The experimentally observed substitution rates for the pool synthesized by electrochemical synthesis (blue) and material deposition-based synthesis (red) as a function of PCR cycles (see main text for details) extrapolate to a base substitution rate which is higher than can be explained by sequencing errors alone.

**Differences in deletion errors between electrochemically-synthesized pools**

Comparing the rate of deletions between the two electrochemically-synthesized oligonucleotide pools shows a 15% increase in mean deletion rate for the GC-constrained pool (considering only the first 102 bases present in both pools, 0.0118 $nt^{-1}$ vs. 0.0136 $nt^{-1}$) compared to the unconstrained pool, see Supplementary Figure 4a. While both pools feature a virtually identical positional dependence of the deletion rate – including the distinct increase in deletion rate after position ~90 – the difference in mean deletion rate indicates that considerable batch-to-batch variation or process changes have affected the fidelity of the electrochemical synthesis.

As expected based on the difference in mean deletion rate, Supplementary Figure 4b and c highlight the differences in the distribution of deletion errors with regards to errors per read and deletion runs between the unconstrained and GC-constrained sample.

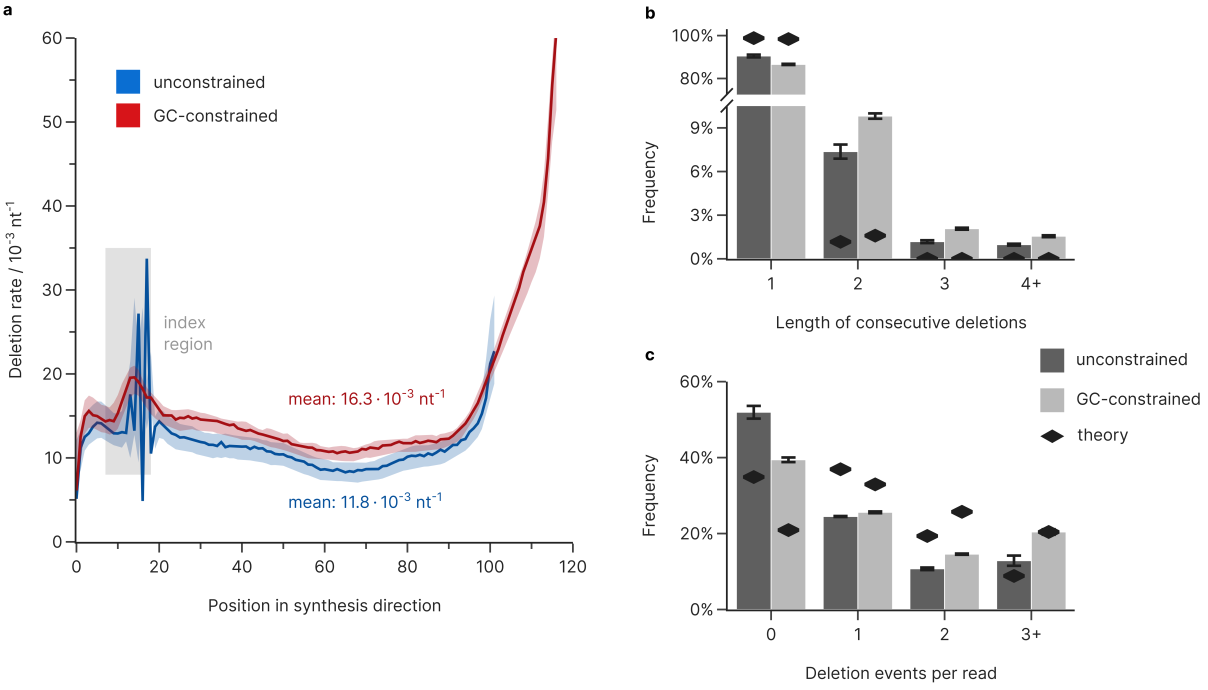


**Supplementary Figure 4: Deletion errors for the two electrochemically-synthesized pools. (a)** Deletion errors as a function of synthesis position for the electrochemically-synthesized pools. The median deletion rate is shown as a function of synthesis position (3'-5' for forward read, 5'-3' for reverse read) for the unconstrained pool (blue, length 102 nt) and the GC-constrained pool (red, length 117 nt). The shaded areas show the rates bounded by the minimum and maximum rates at every position. Co-synthesized priming regions flanking the data-encoding bases are not considered, and the indexing region, where the sequences have very low diversity, is shown as a grey area. **(b)** The

### Supplementary Note 4

**Substitutions during amplification of electrochemically-synthesized pools**

The electrochemically-synthesized oligonucleotide pools were subjected to a different amplification procedure compared to those synthesized by material deposition (see SN X). Instead of consecutive amplifications yielding samples separated by 15 PCR cycles, samples of the electrochemically-synthesized pools were individually amplified with 10 to 30 PCR cycles in a single run, yielding samples separated by only 5 PCR cycles. Adequate dilution of the samples ensured no primer or nucleotide depletion occurred during amplification.

With this procedure, the experimental effort is drastically reduced compared to the sequential amplification, which would allow more experimental conditions to be investigated in the context of errors during amplification (i.e. different polymerases, buffers, additives). However, the procedure has two main downsides:

1) Due to the exponential nature of PCR, there is an upper limit on the range of PCR cycles that can be performed in a single run. At higher numbers of PCR cycles, resource depletion becomes an issue, while at the same time the required dilution would reduce mean oligo coverage drastically below 1.
2) The expected increase in error rates over the limited range of only ~20 cycles is low, especially for higher-fidelity polymerases. This reduces accuracy of the analysis.

Supplementary Figure 5 shows the analysis of substitution errors for the electrochemically-synthesized pools using this procedure. While the overall substitution bias is almost identical, the total substitution rate estimated from this procedure is $1.97 \cdot 10^{-4}$ $nt^{-1}$ $cycle^{-1}$, compared to $1.09 \cdot 10^{-4}$ $nt^{-1}$ $cycle^{-1}$ for the sequential amplification using the pools synthesized by material deposition. Both estimates are within the range estimated in literature for Taq-based polymerases ($1 \cdot 10^{-5}$ to $2 \cdot 10^{-4}$ $nt^{-1}$ $cycle^{-1}$).[8–10] The discrepancy in mean substitution rate is likely the result of the aforementioned limitations of the amplification procedure, as evidenced by the high variation of replicates compared to the equivalent analysis in the main text. Moreover, temperature-induced cytosine deamination rather than polymerase-mediated errors could partly account for the increase in substitution rate,[8] due to the increased duration of thermocycling (up to 30 PCR cycles) compared to the sequential approach (15 cycles per run).

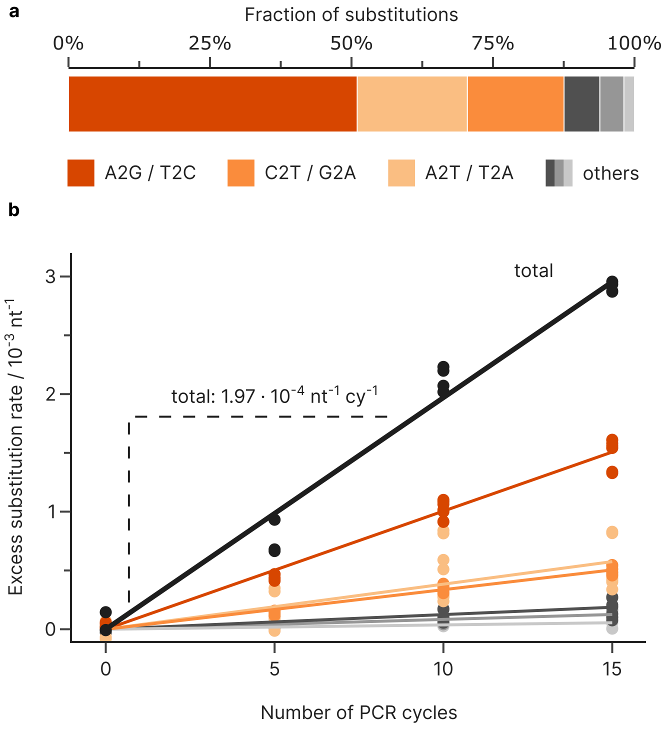


**Supplementary Figure 5: Substitution errors during amplification of the electrochemically-synthesized pools.** Substitutions introduced as a function of the number of additional PCR cycles for the oligonucleotide pools from electrochemical synthesis. **(a)** Substitutions are preferentially introduced as A2G/T2C (51.0%), C2T/G2A (17.1%), and A2T/T2A (19.5%) mutations. **(b)** The regression lines of substation rates as a function of PCR cycles, using the substitution rate at the minimum cycle count as the baseline, show a clear linear trend, equivalent to a total substitution rate of $1.97{\cdot}10^{-4}$ $nt^{-1}$ $cy^{-1}$.

### Supplementary Note 5

**Internal validation of the model for the DNA data storage process**

For the implementation of the experimental conditions, the following error rates and biases were used:

- Material deposition-based synthesis was modelled using the error rates and biases discussed in the main text, including a lognormal coverage distribution, as well as position-, length-, and base-dependent deletion errors.
- Electrochemical synthesis used the specific error rates and coverage biases depending on GC-constraint, as shown in Supplementary Note 3 and the main text. This includes a lognormal coverage distribution, as well as position-, length-, and base-dependent deletion errors.
- Amplification by PCR used the mean substitution rates and base biases estimated from the pools by electrochemical or material deposition-based synthesis, respectively (see Supplementary Note 4 and the main text). The bias in the efficiency distribution used the standard deviation estimated from the amplification experiments of the unconstrained pool synthesized by material deposition (see main text, $\sigma = 0.0051$).
- Aging was modelled using the mean substitution rate and base bias shown in the main text. The extent of aging for each sample was quantified via the number of half-life events (see Supplementary Table 5). No influence on the coverage bias was included, in accordance with the analysis in the main text.
- Sequencing was modelled with 1 000 000 paired reads, including the position-, read-, and base-dependent substitution rates estimated from the PhiX-based error analysis (see main text).
- Dilutions were performed identically as in the experimental protocol.

The scripts for the simulation of all experimental conditions are provided with the code in the repository for a complete documentation of the used parameters.

Supplementary Figure 6 shows a direct comparison of the overall error rates, coverage biases, and the substitution patterns between the experiments and the model output. Coverage bias is quantified by the standard deviation of the logarithmic normalized coverage distribution, in accordance with the lognormal distribution observed for the coverage distribution in the experiments (see main text). Agreement between modelled and experimental error rates and biases is good, with most estimates within 20% of the true value.

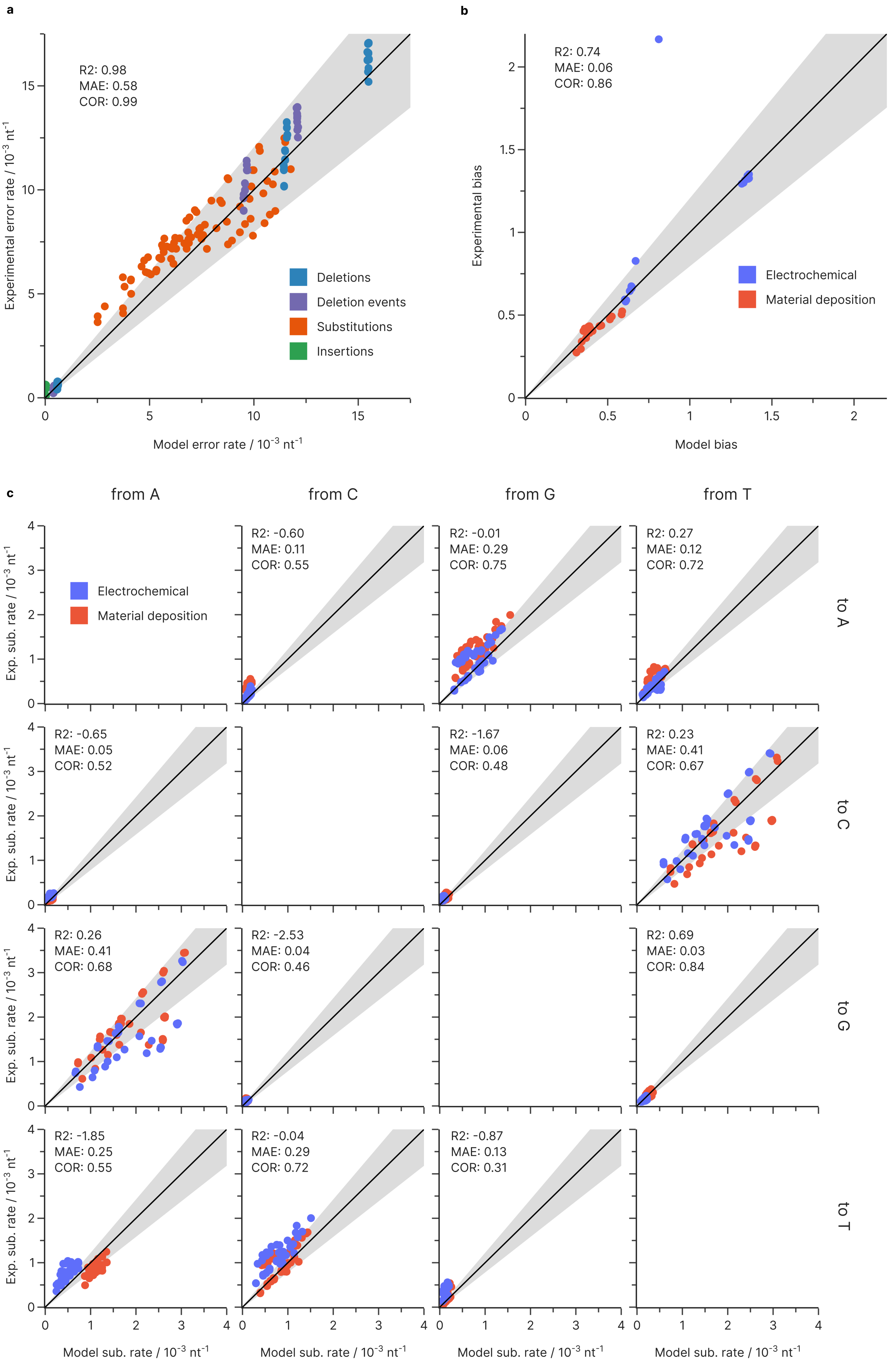
a
R2: 0.98
MAE: 0.58
COR: 0.99
Deletions
Deletion events
Substitutions
Insertions
Experimental error rate / 10-3 nt-1
Model error rate / 10-3 nt-1
b
R2: 0.74
MAE: 0.06
COR: 0.86
Electrochemical
Material deposition
Experimental bias
Model bias
c
from A
from C
from G
from T
to A
to C
to G
to T
Electrochemical
Material deposition
R2: -0.60
MAE: 0.11
COR: 0.55
R2: -0.01
MAE: 0.29
COR: 0.75
R2: 0.27
MAE: 0.12
COR: 0.72
R2: -0.65
MAE: 0.05
COR: 0.52
R2: -1.67
MAE: 0.06
COR: 0.48
R2: 0.23
MAE: 0.41
COR: 0.67
R2: 0.26
MAE: 0.41
COR: 0.68
R2: -2.53
MAE: 0.04
COR: 0.46
R2: 0.69
MAE: 0.03
COR: 0.84
R2: -1.85
MAE: 0.25
COR: 0.55
R2: -0.04
MAE: 0.29
COR: 0.72
R2: -0.87
MAE: 0.13
COR: 0.31
Exp. sub. rate / 10-3 nt-1
Model sub. rate / 10-3 nt-1

**Supplementary Figure 6: Internal validation of error rates and coverage biases. (a)** Comparison of mean rates of deletions (blue), deletion events (purple), substitutions (red), and insertions (green) between experiments and simulation for all conditions in this study. The black line denotes the diagonal, and the shaded area denotes a deviation of ±20%. **(b)** Comparison of coverage bias, quantified by the standard deviation of the lognormal distribution of reads, between experiments and simulation for all conditions in this study, grouped by conditions using electrochemical (blue) or material deposition-based synthesis (red). The black line denotes the diagonal, and the shaded area denotes a deviation of ±20%. **(c)** Direct comparison of the substitution rates, grouped by electrochemical (blue) or material deposition-based synthesis (red), and for each substitution type, between experiments and simulation for all conditions in this study. The black line denotes the diagonal, and the shaded area denotes a deviation of ±20%.

For the overall error rates, the largest deviation is observed for the substitution rate of samples with few PCR cycles. For these experimental conditions, the model underestimates the substitution rate by a constant offset of around 1-2·$10^{-3}$ $nt^{-1}$. It is likely that this deviation is caused by the aforementioned unexplained source of substitutions attributed to synthesis errors (see Supplementary Note 3 and Supplementary Figure 3), which was estimated at around 0.8·$10^{-3}$ $nt^{-1}$. Moreover, all error types show a high variability relative to the value of the error rate, likely caused by the statistical nature of the experiments and the analysis procedure.

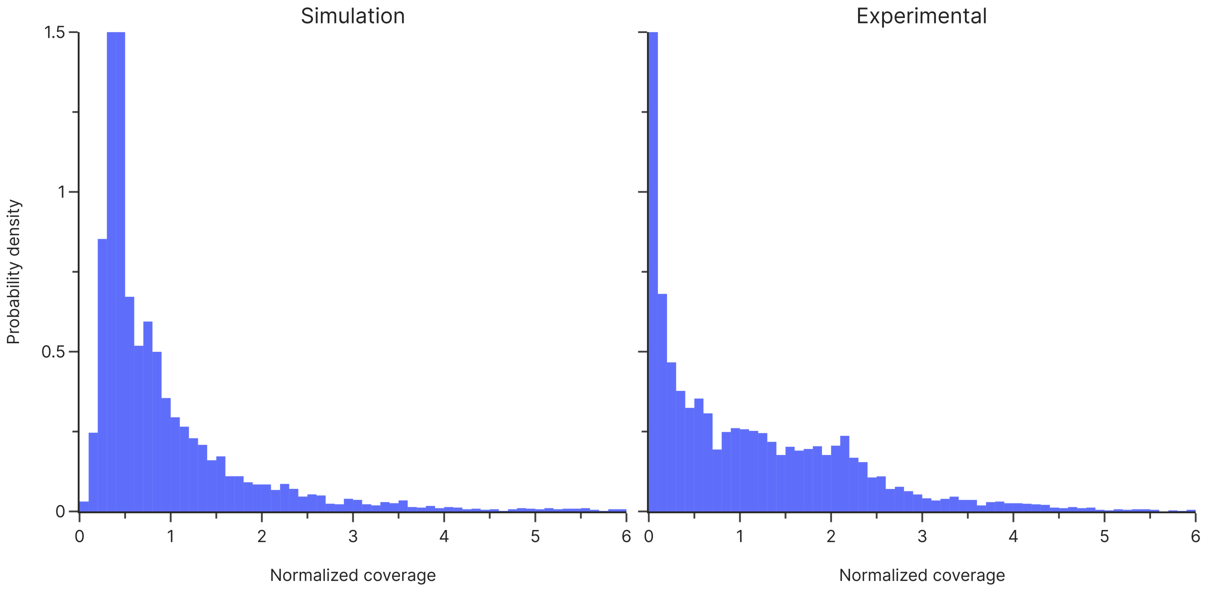


**Supplementary Figure 7: Comparison of the normalized coverage for the last sample of the amplification experiment for the unconstrained, electrochemically-synthesized pool.** The large dilution factor in both the experiment and the simulation leads to a long-tailed distribution of normalized oligo coverages.

The comparison of the coverage bias shows good agreement both for the initial bias, as well as its evolution during amplification. The final amplification experiment of the unconstrained, electrochemically-synthesized pool (30 cycles) is the major exception, for which the coverage bias is underestimated considerably. As described in Supplementary Note 4, this is caused by the extreme dilution required to facilitate amplification without resource depletion for so many cycles, leading to a mean coverage of only 0.43 oligos per sequence. As a result, coverage bias is highly dependent on this stochastic dilution event and its estimation via the standard deviation of the corresponding lognormal distribution does not accurately show the accuracy of prediction. Supplementary Figure 7 compares the full coverage distribution of this sample with the model output, highlighting the difference in skew of the distribution. Importantly, the sequence dropout predicted by the model (52%) is comparable to the experimental result (53%).

The agreement between model and experiment for the substitution patterns (see Supplementary Figure 6c) is mixed. While all substitution patterns with a large error rate (e.g. A2G, C2T, G2A, T2C) are reasonably well reproduced by the model, as highlighted by the correlation coefficients given in Supplementary Figure 6c, some other substitution patterns (i.e. A2T, C2A, G2T) are not. Some of this deviation can be explained by the aforementioned, unexplained source of substitutions attributed to synthesis errors (see Supplementary Note 4 and Supplementary Figure 3), which was estimated at around $0.8 \cdot 10^{-3}$ $nt^{-1}$. This hypothesis would explain some of the observed systematic deviation in base-specific substitution rates between the two synthesis providers (e.g. clearly visible for A2T and C2A substitutions). Still, the magnitude of the unexplained substitution rate is negligibly small, at less than half that caused by sequencing, such that its omission is irrelevant to practical applications.

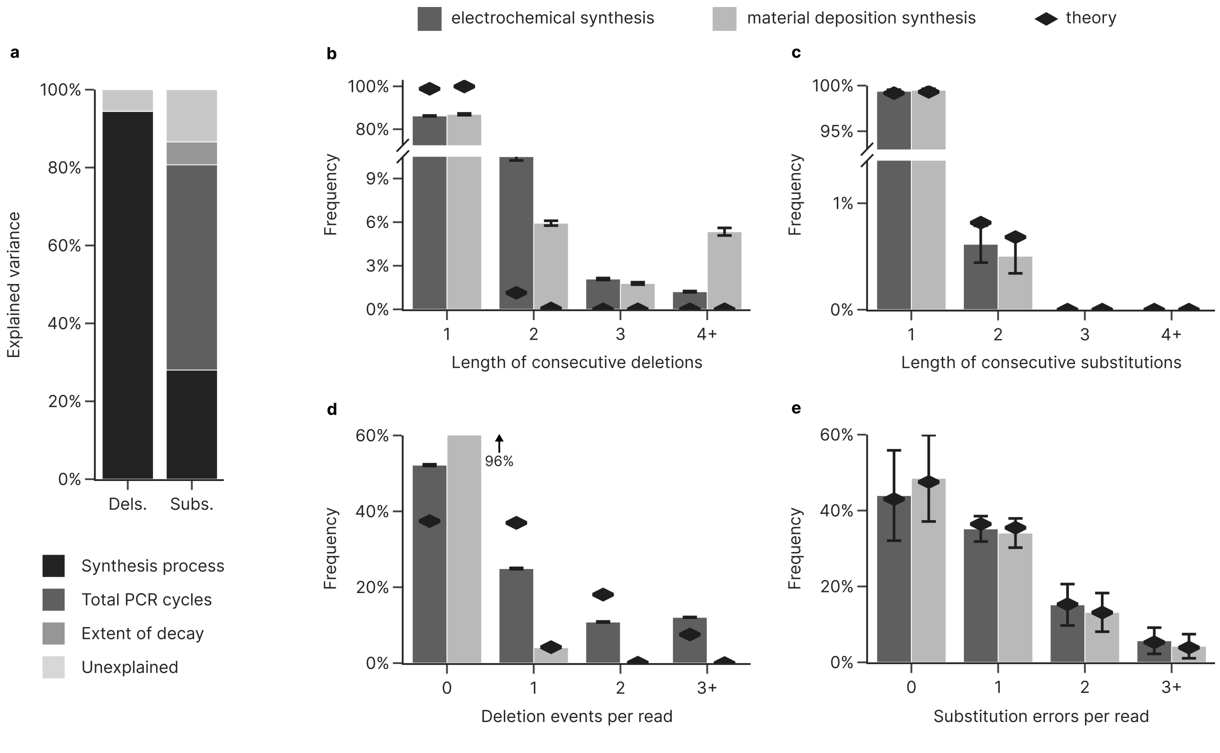


**Supplementary Figure 8: Validation of error variance and independence.** The analysis is identical to Figure 1 in the main text, but uses the simulated sequencing data instead. (**a**) The contribution of synthesis process, presence of GC-constraint, and sample preparation to the overall variance in mean deletion (left) and substitution (right) rates between samples was assessed by four-way analysis of variance (ANOVA). (**b-e**) Distributional analysis of error independence for deletions (b+d) and substitutions (c+e) based on the observed frequency of error runs (b+c) and errors per read (d+e), for the GC-unrestricted pools synthesized by electrochemical (dark grey) and material deposition (light grey) processes. Theoretical distributions under the assumption of error independence are also shown (black diamonds, geometric/binomial). The histogram for deletions per read treats any run of deletions as a single event to accommodate the non-ideality of deletion runs. Error bars show the standard deviation of the sample.

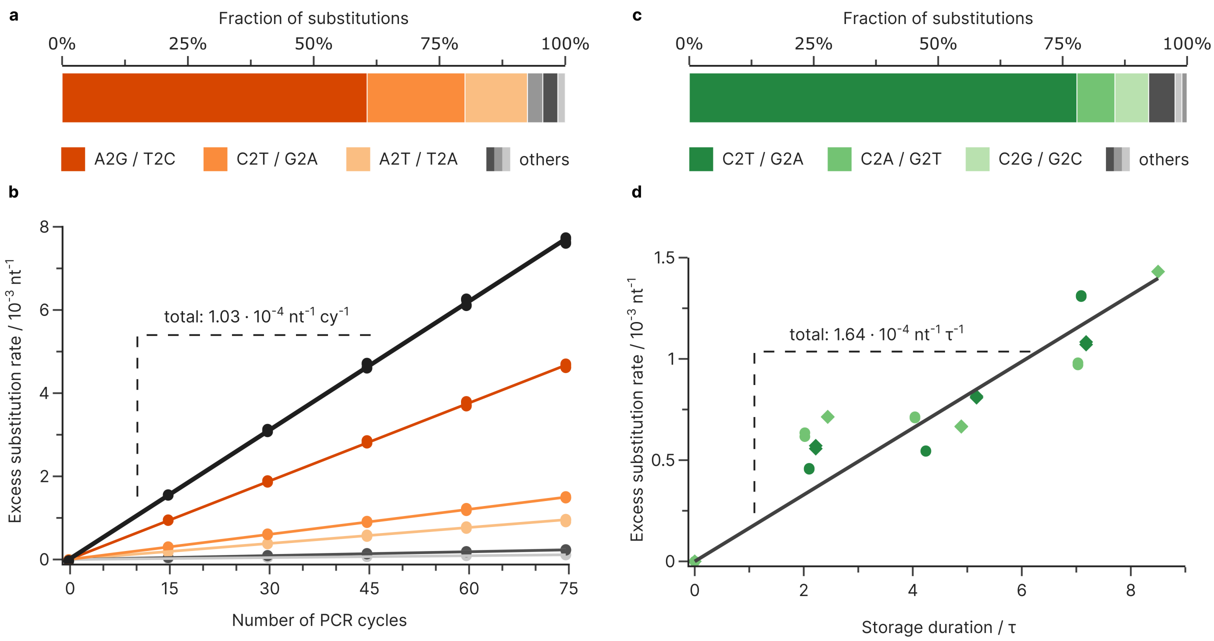


**Supplementary Figure 9: Validation of errors introduced during amplification and storage.** The analysis is identical to Figures 3 and 4 in the main text, but uses the simulated sequencing data instead. (**a+b**) Substitutions introduced as a function of the number of additional PCR cycles for the oligonucleotide pools from material deposition-based synthesis, using the substitution rate at 15 cycles as the baseline. The regression slope (solid lines) yields an overall error rate of $1.03{\cdot}10^{-4}$ $nt^{-1}$ per cycle and shows A2G/T2C transitions account for 61% of substitutions, followed by C2T/G2A transitions (20%) and A2T/T2A transversions (13%). (**b+c**) Substitutions introduced as a function of the total storage duration in half-lives, using the error rates of the unaged reference as baseline. Substitutions increase at a rate of $1.56{\cdot}10^{-4}$ $nt^{-1}$ per half-live based on the regression slope (solid line). Substitutions are mainly C2T/G2A transitions (dark green, 77%) with minor C2A/G2T and C2G/G2C transversions (7% and 6% respectively).

To validate the correct reproduction of the (partially non-ideal) distribution of errors per read and error runs, we reproduced the validation of error independence for the simulated datasets in Supplementary Figure 6. As desired, the distribution of consecutive deletions and deletion events per read accurately follows the non-ideal distribution observed experimentally, while substitutions behave independently, as expected under the assumption of error independence (compare to main text). Similarly, the experimentally observed mean substitution rates and base preferences during amplification and aging are accurately reproduced by the model, as shown in Supplementary Figure 9.

#### External validation of the model for the DNA data storage process

To compare the errors and biases introduced by the process model of the digital twin against an external reference, the generational experiments performed by Koch et al.[4] were recreated with our tool and underwent identical post-processing and error analysis. For the implementation of the experimental conditions, the following parameters were used:

- The oligonucleotide used by Koch et al. were obtained from CustomArray, thus synthesis was modelled as electrochemical synthesis using the error rates and coverage biases estimated from our experimental data, as presented in the main text (i.e. not using the specific deletion patterns described for unconstrained and GC-constrained pools in Supplementary Note 3). Due to the GC-constraint imposed by their encoder, the design sequences were limited to a GC content of 45-55%. Since this is broader than the GC content of our GC-constrained datasets (only 50%) and narrower than the unconstrained datasets (30-70%), we used the arithmetic mean of the GC-constrained and unconstrained coverage biases (i.e. $\sigma = 0.94$) as the initial bias for the pool of the generational experiments.

- For amplification by PCR, Koch et al. also used KAPA SYBR FAST qPCR Master Mix, so a polymerase fidelity of 1 and the substitution patterns as analysed in the main text were assumed. The standard deviation of the normalized efficiency distribution was chosen as $\sigma = 0.012$, based on the analysis of the data presented in the main text.
- In accordance with the experimental protocol by Koch et al., no aging was included in the model.
- Sequencing was assumed identical to the error rates estimated from our PhiX data, identically to the internal validation described above.
- Insufficient information about the exact dilutions and concentrations used throughout the workflow is presented in the study by Koch et al. Thus, most dilutions were estimated from the reported number of PCR cycles needed for amplification. However, this should not influence model estimations significantly, as no dilution leads to a low mean oligo coverage for which relevant stochastic effects are expected.

A flowsheet describing the assumed workflow is shown in Supplementary Figure 26. The scripts for the simulation of all generational experiments from Koch et al.[4] are provided with the code in the repository for a complete documentation of the used parameters.

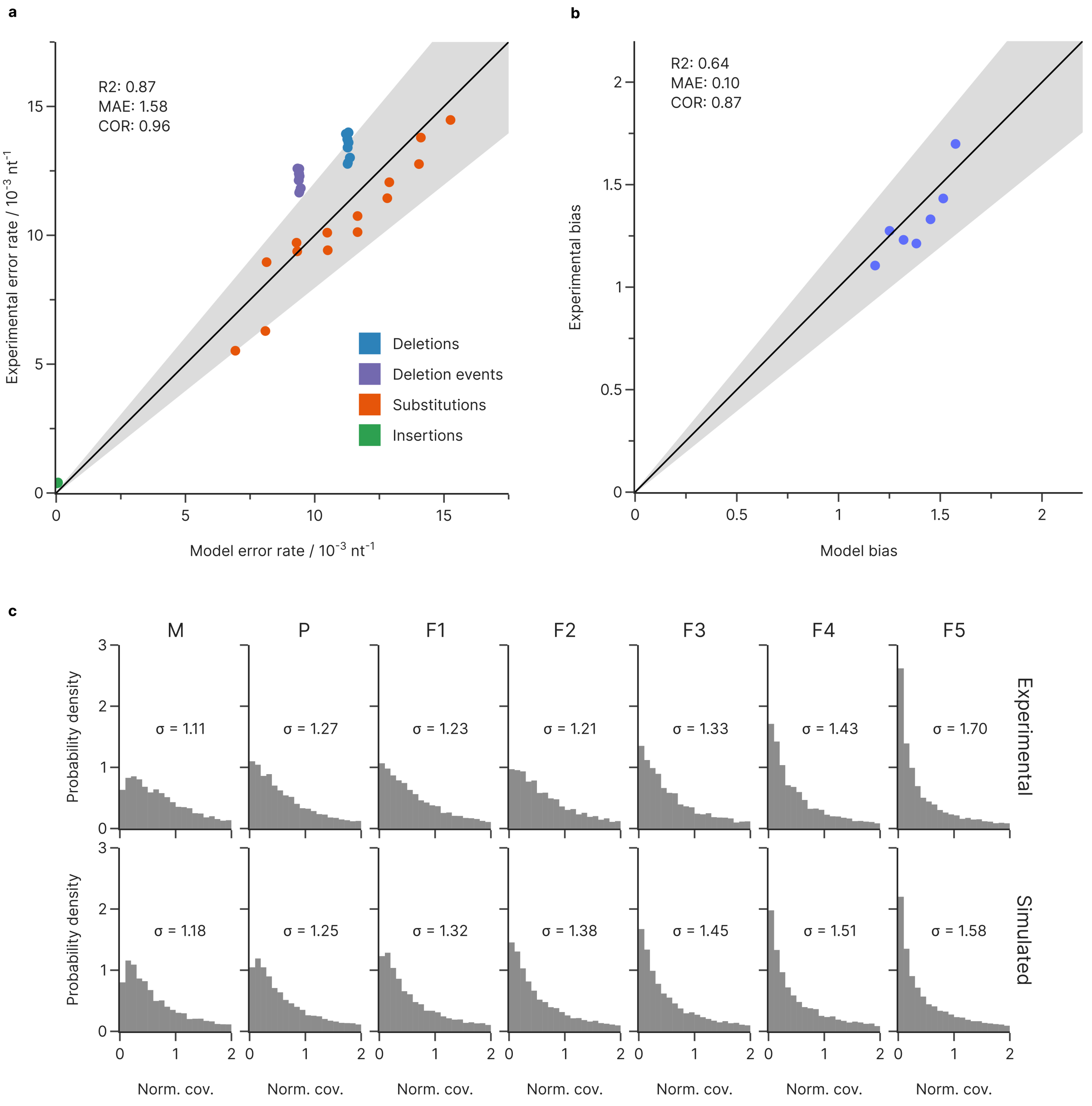


**Supplementary Figure 10: External validation of error rates and coverage biases. (a)** Comparison of mean rates of deletions (blue), deletion events (purple), substitutions (red), and insertions (green) between the generational experiments by Koch et al.[4] and the simulation. The black line denotes the diagonal, and the shaded area denotes a deviation of ±20%. **(b)** Comparison of coverage bias, quantified by the standard deviation of the lognormal distribution of reads, between the generational experiments by Koch et al.[4] and the simulation. The black line denotes the diagonal, and the shaded area denotes a deviation of ±20%. **(c)** Direct comparison of the coverage distributions obtained experimentally and from the simulation for all generations.

### Supplementary Tables

**Supplementary Table 1: Design of oligonucleotide pools.** The sequences for all pools were designed according to the workflow described by Meiser et. al.[11] All sequences include priming regions at the beginning and end (20+21 nt), which are denoted as +41 in the sequence length. Pools with fixed GC feature a GC-constraint that only allows sequences with 50% GC content. Generally, all sequences are pseudo-random and thus feature a homogeneous base content, with the exception if the unconstrained pool from Genscript, which includes an index region which is heterogeneous. Design sequences are provided with the sequencing data in the repository.

| Synthesis | | Design sequences | | Comments |
|---|---|---|---|---|
| Provider | GC | Number | Length | |
| Genscript | all | 12472 | 102+41 | previously used in Ref. 11, index present |
| Genscript | fixed | 12402 | 117+41 | random sequences, no index |
| Twist | all | 12000 | 108+41 | random sequences, no index |
| Twist | fixed | 12000 | 108+41 | random sequences, no index |

**Supplementary Table 2: Primer sequences.** Underlined bases highlight the index region for Illumina multiplexing.

| Name | Sequence (5'-3') |
|---|---|
| **0F** | ACACGACGCTCTTCCGATCT |
| **0R** | AGACGTGTGCTCTTCCGATCT |
| **2FUF** | AATGATACGGCGACCACCGAGATCTACACTCTTTCCCTACACGACGCTCTTCCGATCT |
| **2RIF-GM5** | CAAGCAGAAGACGGCATACGAGATCACTGTGTGACTGGAGTTCAGACGTGTGCTCTTCCGATCT |
| **2RIF-GM7** | CAAGCAGAAGACGGCATACGAGATGATCTGGTGACTGGAGTTCAGACGTGTGCTCTTCCGATCT |
| **2RIF-GM8** | CAAGCAGAAGACGGCATACGAGATTCAAGTGTGACTGGAGTTCAGACGTGTGCTCTTCCGATCT |
| **2RIF-GM10** | CAAGCAGAAGACGGCATACGAGATAAGCTAGTGACTGGAGTTCAGACGTGTGCTCTTCCGATCT |
| **2RIF-GM11** | CAAGCAGAAGACGGCATACGAGATGTAGCCGTGACTGGAGTTCAGACGTGTGCTCTTCCGATCT |
| **2RIF-GM12** | CAAGCAGAAGACGGCATACGAGATTACAAGGTGACTGGAGTTCAGACGTGTGCTCTTCCGATCT |
| **2RIF-GM17** | CAAGCAGAAGACGGCATACGAGATCTCTACGTGACTGGAGTTCAGACGTGTGCTCTTCCGATCT |

**Supplementary Table 3: Parameters of the calibration curves for qPCR analysis of the oligonucleotide pools.** Parameter estimates are given with their standard error.

| Sample | Slope | Intercept | R2 | Efficiency |
|---|---|---|---|---|
| **Genscript (unconstrained)** | -3.493±0.029 | 0.81±0.15 | 0.9993 | 93.3% |
| **Genscript (GC-constrained)** | -3.465±0.009 | 1.90±0.03 | 0.9999 | 94.3% |
| **Twist (unconstrained)** | -3.660±0.021 | 1.44±0.11 | 0.9995 | 87.6% |
| **Twist (GC-constrained)** | -3.713±0.017 | 1.41±0.09 | 0.9997 | 85.9% |

**Supplementary Table 4: Parameters of the decay model for aging of the oligonucleotide pools.** Parameter estimates are given with their standard error.

| Sample | $k$ / $d^{-1}$ | $\tau$ / d | $R^2$ |
|---|---|---|---|
| **Genscript (unconstrained)** | 0.74±0.07 | 0.93±0.09 | 0.971 |
| **Genscript (GC-constrained)** | 0.66±0.07 | 0.96±0.09 | 0.976 |
| **Twist (unconstrained)** | 0.70±0.11 | 1.00±0.16 | 0.925 |
| **Twist (GC-constrained)** | 0.84±0.12 | 0.82±0.12 | 0.944 |

**Supplementary Table 5: Conversion of storage duration to number of half-lives for all oligonucleotide pools.** Parameter estimates are given with their standard deviation.

| Sample | Time / d | Rel. conc. | # Half-lives |
|---|---|---|---|
| **Genscript (unconstrained)** | 0.00 | 1.00±0.19 | 0.00 |
| | 1.95 | 0.113±0.013 | 2.10 |
| | 3.95 | 0.105±0.101 | 4.24 |
| | 6.61 | 0.0061±0.0028 | 7.09 |
| **Genscript (GC-constrained)** | 0.00 | 1.00±0.04 | 0.00 |
| | 2.13 | 0.083±0.017 | 2.22 |
| | 4.97 | 0.027±0.006 | 5.17 |
| | 6.91 | 0.0094±0.0009 | 7.18 |
| **Twist (unconstrained)** | 0.00 | 1.00±0.28 | 0.00 |
| | 2.01 | 0.074±0.008 | 2.02 |
| | 4.03 | 0.028±0.012 | 4.04 |
| | 7.00 | 0.017±0.005 | 7.03 |
| **Twist (GC-constrained)** | 0.00 | 1.00±0.02 | 0.00 |
| | 2.01 | 0.035±0.024 | 2.44 |
| | 4.03 | 0.044±0.041 | 4.89 |
| | 7.00 | 0.0038±0.0018 | 8.50 |

**Supplementary Table 6: Distribution of deletion errors based on deleted base for both synthesis processes.** Parameter estimates are given with their standard deviation.

| Deleted base | Electrochemical | | Material deposition | |
|---|---|---|---|---|
| | Forward read | Reverse read | Forward read | Reverse read |
| **A** | 27.1±3.2% | 24.6±1.3% | 24.7±0.4% | 25.1±0.4% |
| **C** | 25.2±0.3% | 23.2±1.5% | 23.6±0.4% | 26.2±0.5% |
| **G** | 23.2±1.6% | 25.2±0.3% | 26.7±0.6% | 23.6±0.6% |
| **T** | 24.5±1.3% | 27.1±3.2% | 25.0±0.7% | 25.1±0.6% |

**Supplementary Table 7: Estimation of the initial abundance of the oligonucleotide pools.** Estimates of the cycle thresholds are given with their standard deviation, based on a plate of 96 wells. An estimate of the initial concentration of amplifiable DNA in all oligonucleotide pools is possible based on the qPCR calibration and the result of pool amplification. Considering also that the pool is first converted from single- to double-stranded DNA (equivalent to a factor of 2), the initial abundance can be directly calculated from the cycle threshold from the calibration curves.

| Sample | Cycle threshold | Dilution factor | Init. conc. / ng $\mu L^{-1}$ |
|---|---|---|---|
| **Genscript (unconstrained)** | 15.97±0.14 | 250 | 0.023 |
| **Genscript (GC-constrained)** | 16.52±0.08 | 250 | 0.030 |
| **Twist (unconstrained)** | 16.07±0.03 | 5000 | 1.01 |
| **Twist (GC-constrained)** | 15.50±0.03 | 5000 | 1.61 |

**Supplementary Table 8: Results of the main-effects, three-way ANOVA for overall error rates (n=80).** The factors synthesis provider, number of PCR cycles, and days of storage were analysed for explaining the variance observed in the mean deletion, substitution, and insertion rates for both forward and reverse reads of all 40 experimental conditions. Type II sum of squares and HC3-correction were used.

| Error type | Factor | SoS | DoF | F | p(>F) | $\eta^2$ |
|---|---|---|---|---|---|---|
| **Deletions** | C(synthesis) | 2.4E-3 | 1 | 933.7 | 1.9E-44 | 92.3% |
| | # PCR cycles | 4.6E-6 | 1 | 1.8 | 0.19 | 0.2% |
| | # days of storage | 3.3E-13 | 1 | 0.0 | 0.99 | 0.0% |
| | Residual | 1.9E-4 | 76 | - | - | 7.5% |
| **Substitutions** | C(synthesis) | 5.1E-5 | 1 | 115.4 | 6.7E-17 | 7.9% |
| | # PCR cycles | 5.6E-4 | 1 | 1251.4 | 5.9E-49 | 86.2% |
| | # days of storage | 0.4E-6 | 1 | 9.38 | 3.0E-3 | 0.6% |
| | Residual | 3.4E-5 | 76 | - | - | 5.2% |
| **Insertions** | C(synthesis) | 3.2E-6 | 1 | 832.8 | 1.1E-42 | 91.4% |
| | # PCR cycles | 8.2E-9 | 1 | 2.1 | 0.15 | 0.2% |
| | # days of storage | 8.5E-14 | 1 | 0.0 | 0.99 | 0.0% |
| | Residual | 2.9E-7 | 76 | - | - | 8.3% |

### Supplementary Figures

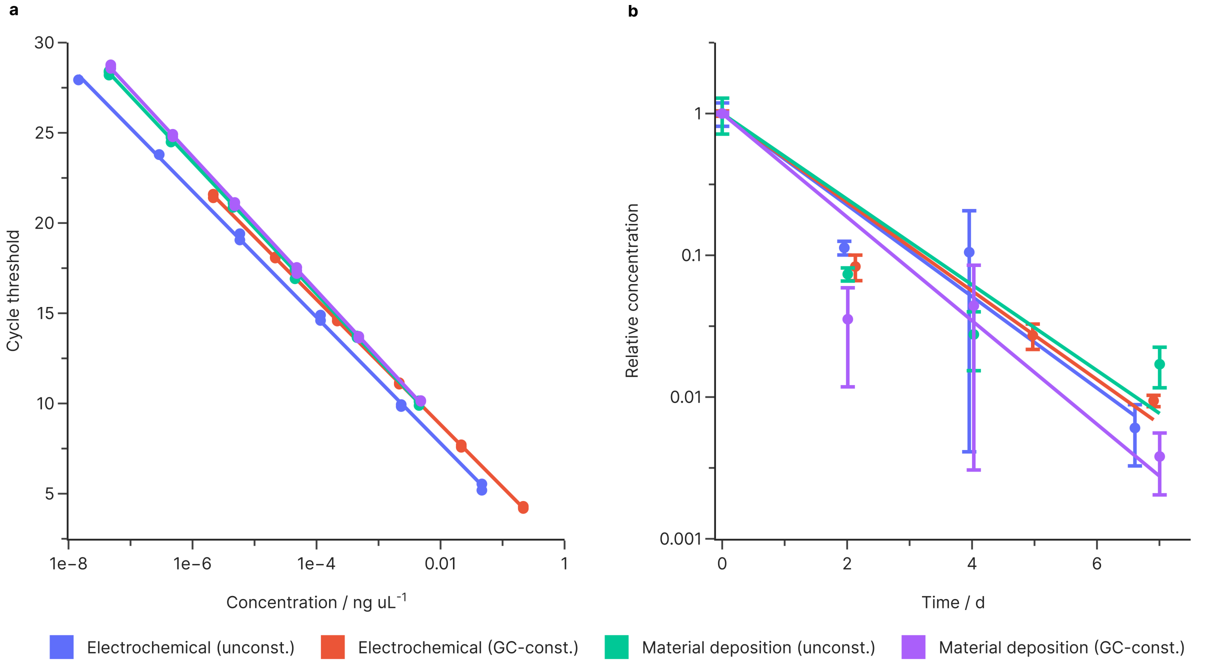


**Supplementary Figure 11: qPCR calibration curve and decay curve. (a)** Calibration was performed by serial dilution of the stock solutions of each pool. Prior quantification was performed using fluorescence on a Qubit spectrophotometer with the Qubit HS DNA kit. The parameters of the regressed calibration curves (solid lines) are given in Supplementary Table 3. **(b)** Decay curves for all oligo pools. By normalizing the measured concentration after aging for a specific duration to the unaged reference, a first-order decay model was fit to the data (solid lines). The parameters of the decay model are given in Supplementary Table 4.

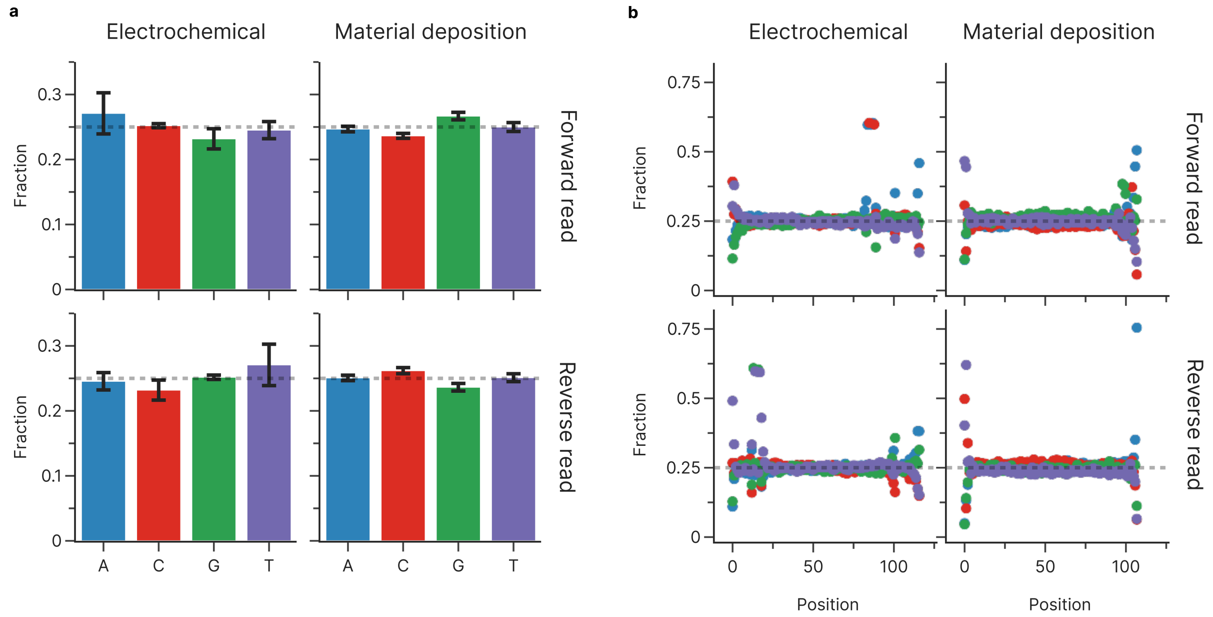


**Supplementary Figure 12: Bias in deletion errors by base and position. (a)** Fraction of deletions by base, grouped by synthesis process and read. Error bars show the standard deviation. Minor bias in the base preference for deletions was observed for both synthesis processes. A one-way ANOVA (Type II, HC3) showed that this bias was statistically significant for both electrochemical ($F(3, 64) = 12.05$, $p = 2e^{-6}$, $\eta^2 = 36\%$) and material deposition-based synthesis ($F(3, 84) = 140.6$, $p = 1e^{-32}$, $\eta^2 = 83\%$). However, the effective change in the fraction of deleted bases was minor, deviating at most 2.1 percentage points from the expected share of 25% per base (see Supplementary Table 6). The distribution of deletions is also mirrored by base pairing between forward and reverse reads, as expected due to amplification by PCR. **(b)** Fraction of deletions by base as a function of position in the sequence, grouped by synthesis process and read. The previously mentioned, minor bias in the base preference was also observed to be stable with regards to the base position, except for regions at the start and end of the oligos. Colours are identical to panel (a).

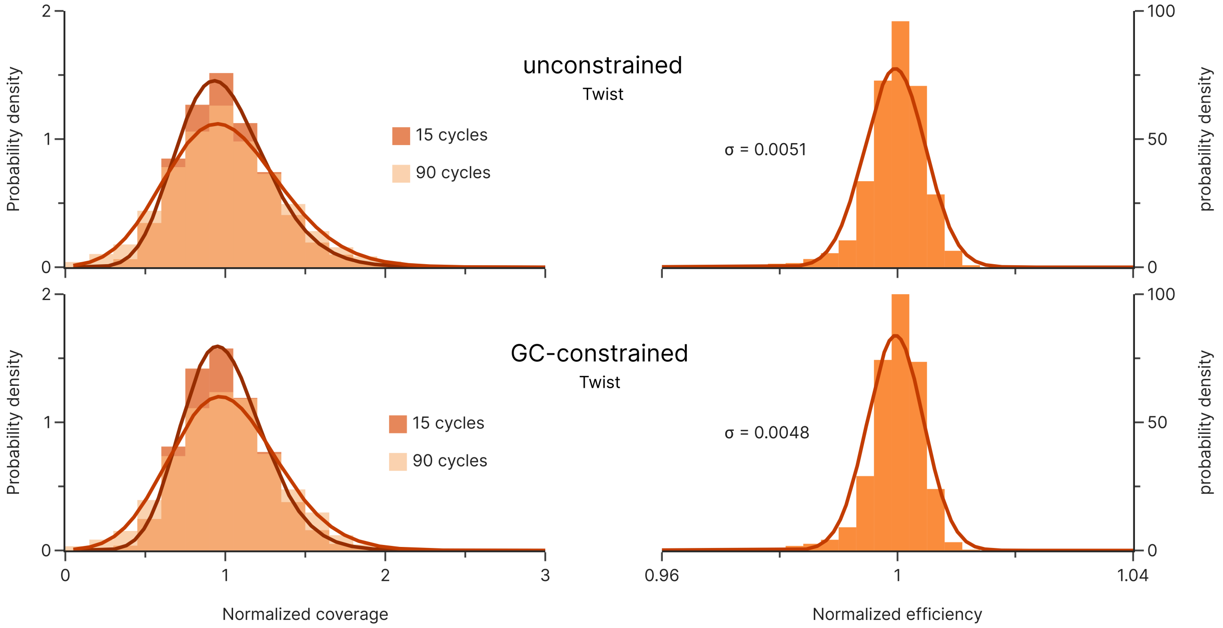


**Supplementary Figure 13: Amplification bias for oligo pools synthesized by material deposition.** The normalized coverage distributions (left) of both unconstrained and GC-constrained sequencing pools before (dark orange) and after repeated amplification (light orange) show near identical broadening of the oligo coverage after PCR. The associated normalized efficiencies (right) also show a similar, narrow distribution. The broadness of the resulting efficiency distribution, characterized by the standard deviation of the fitted normal distributions, is given in the figure.

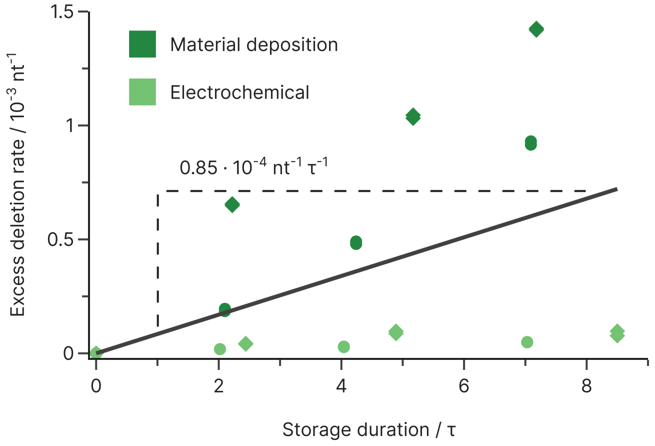


**Supplementary Figure 14: Deletions due to storage.** Deletions introduced as a function of the total storage duration in half-lives, using the error rates of the unaged reference as baseline. Deletions increase at an average rate of $0.83{\cdot}10^{-4}$ $nt^{-1}$ per half-live (solid line). The changes in deletion rate are different between the pools synthesized by material deposition and electrochemical synthesis, hinting that the negligible increase in deletions (e.g. <1% for electrochemically pools) is likely caused by stochastic effects or noise in the error analysis.

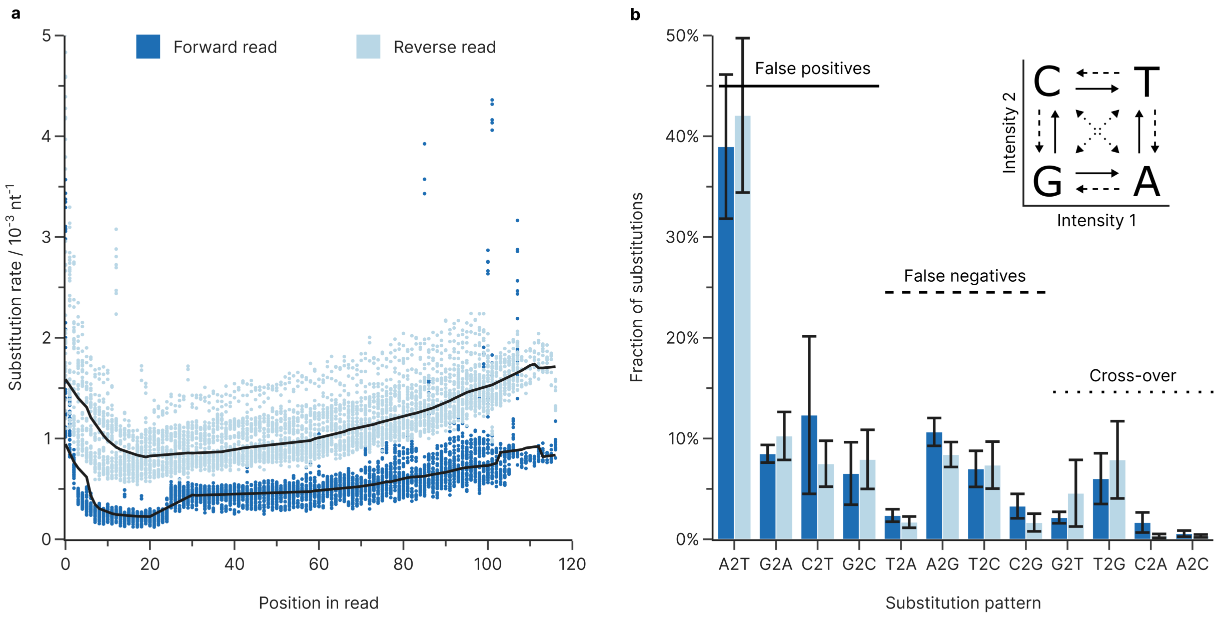


**Supplementary Figure 15: Sequencing errors estimated from paired reads**. The identification of mismatches between paired sequencing reads is also frequently used to quantify sequencing errors, relying on the assumption that the sequencing errors between both paired reads are independent.[12] **(a)** Substitution rate during sequencing as a function of position and read (forward read, dark blue; reverse read, light blue) estimated from paired reads, and **(b)** overall base preference for substitutions during sequencing. Comparing our PhiX-derived sequencing errors to those obtained from this paired-read approach, we found good agreement for the cycle-dependence and base preference of substitution errors. The mean substitution rate estimated from paired reads was lower however, at only $8.5\cdot10^{-4}$ nt$^{-1}$, and A2T transversions were more prevalent, at around 40%. While this discrepancy can be explained by the presence of errors in the PhiX standard, it is also caused by the neglection of clonal amplification in the paired reads-based approach. Errors introduced during the initial cycles of this clonal amplification, while contributing to the overall sequencing error, are present on most reads in a cluster and will thus not be identified by the paired-read approach. This hypothesis is corroborated by the gain in base transitions (e.g. A↔G and C↔T) in the PhiX-derived base bias, which closely matches the bias signature of polymerases,[8,10] especially the 3' exonuclease-deficient polymerases used in SBS.[13] Therefore, our PhiX-based approach likely overestimates the rate of sequencing errors, whereas the paired reads-based approach underestimates them.

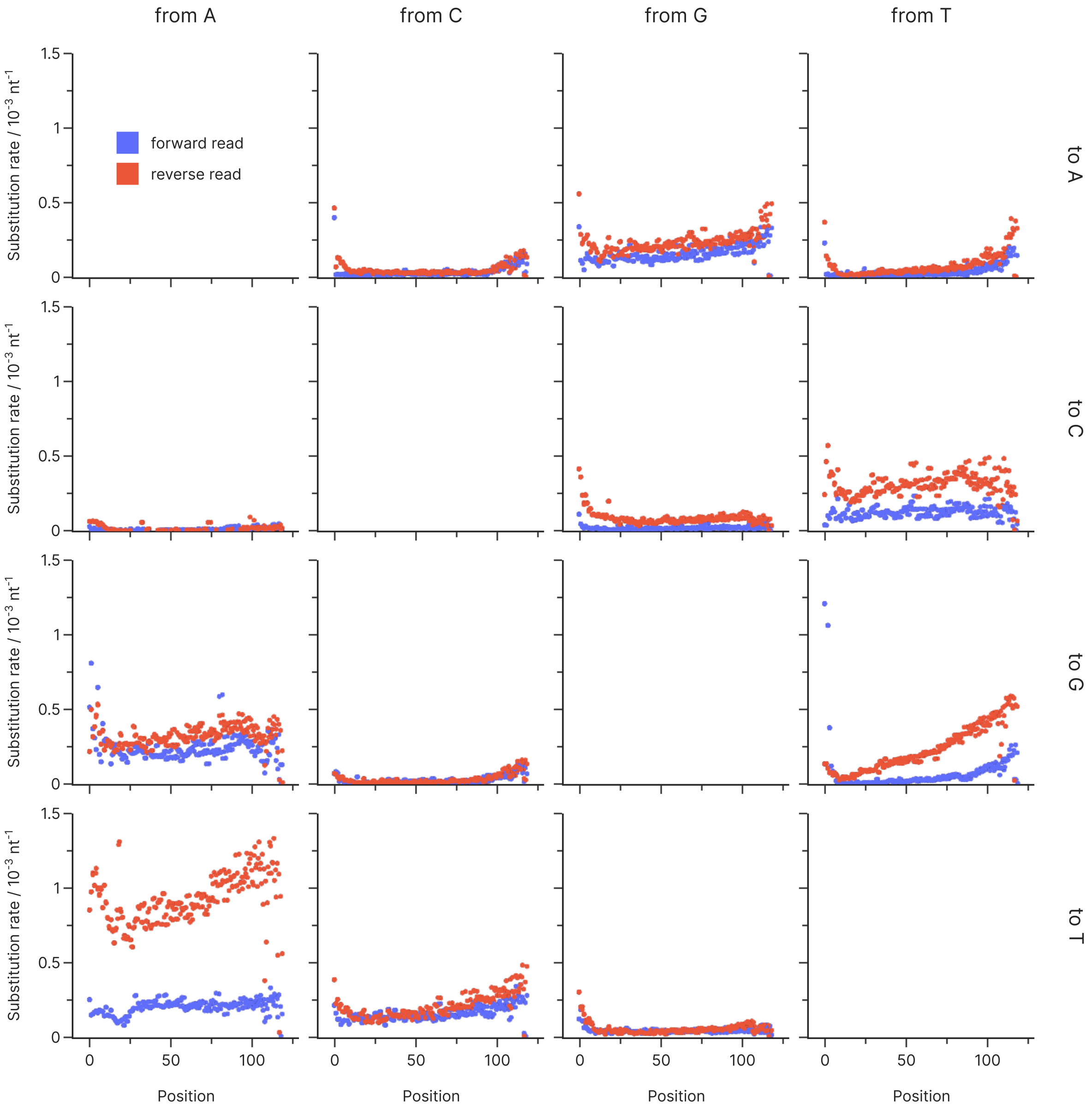


**Supplementary Figure 16: Positional dependence of sequencing errors.** The positional dependence of the PhiX-derived sequencing errors in our dataset explains both the large difference in mean substitution rate between forward and reverse reads and the increase in substitution rate towards the end of both sequencing reads. The large difference between reads is caused primarily by a 3-5x higher rate of A2T transversions in the reverse read, as well as higher rates of T2C and T2G substitutions. While most substitution rates remain approximately constant throughout the sequencing run, the increase in substitution rate towards the end of the reads is dominated by A2T, T2G, C2T, G2A, and T2A substitutions. With the exception of T2G, all of these substitution patterns correspond to a miscall in one of the sequencer's two intensity channels.[14]

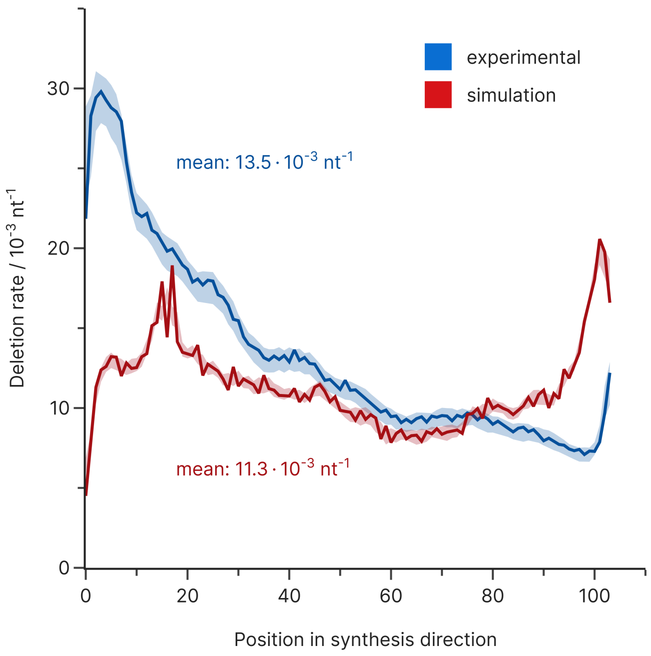


**Supplementary Figure 17: Deletion errors for the generational experiments by Koch et al.** Deletion errors as a function of synthesis position for both the experimental dataset by Koch et al. and the simulation in this study. The median deletion rate is shown as a function of synthesis position (3’-5’ for forward read, 5’-3’ for reverse read) for the experimental dataset (blue, length 102 nt) and the simulated dataset (red, length 117 nt). The shaded areas show the rates bounded by the minimum and maximum rates at every position. Co-synthesized priming regions flanking the data-encoding bases are not considered.

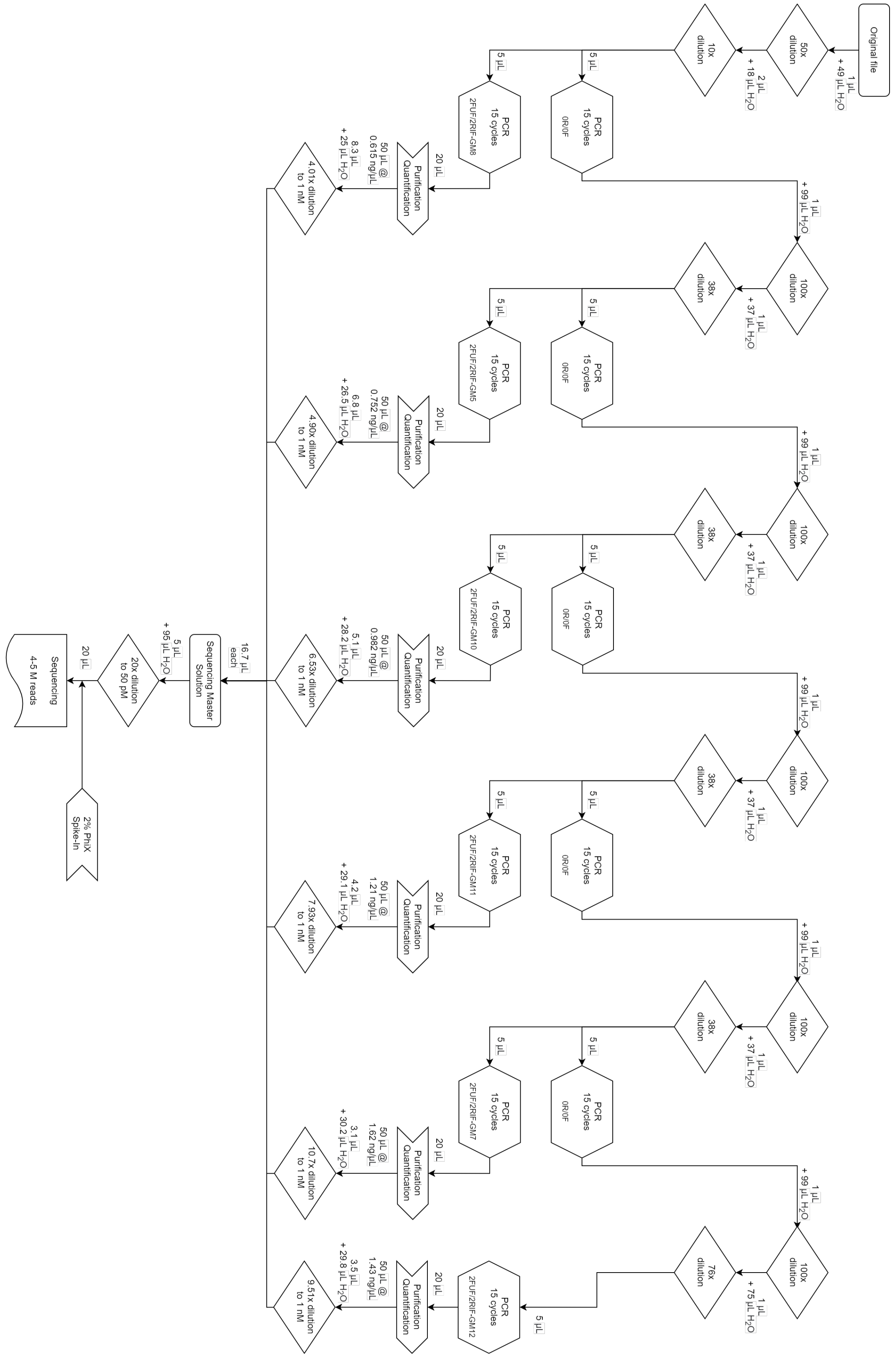


**Supplementary Figure 18: Experimental procedure for the amplification experiments using the unconstrained oligonucleotide pool synthesized by material deposition.**

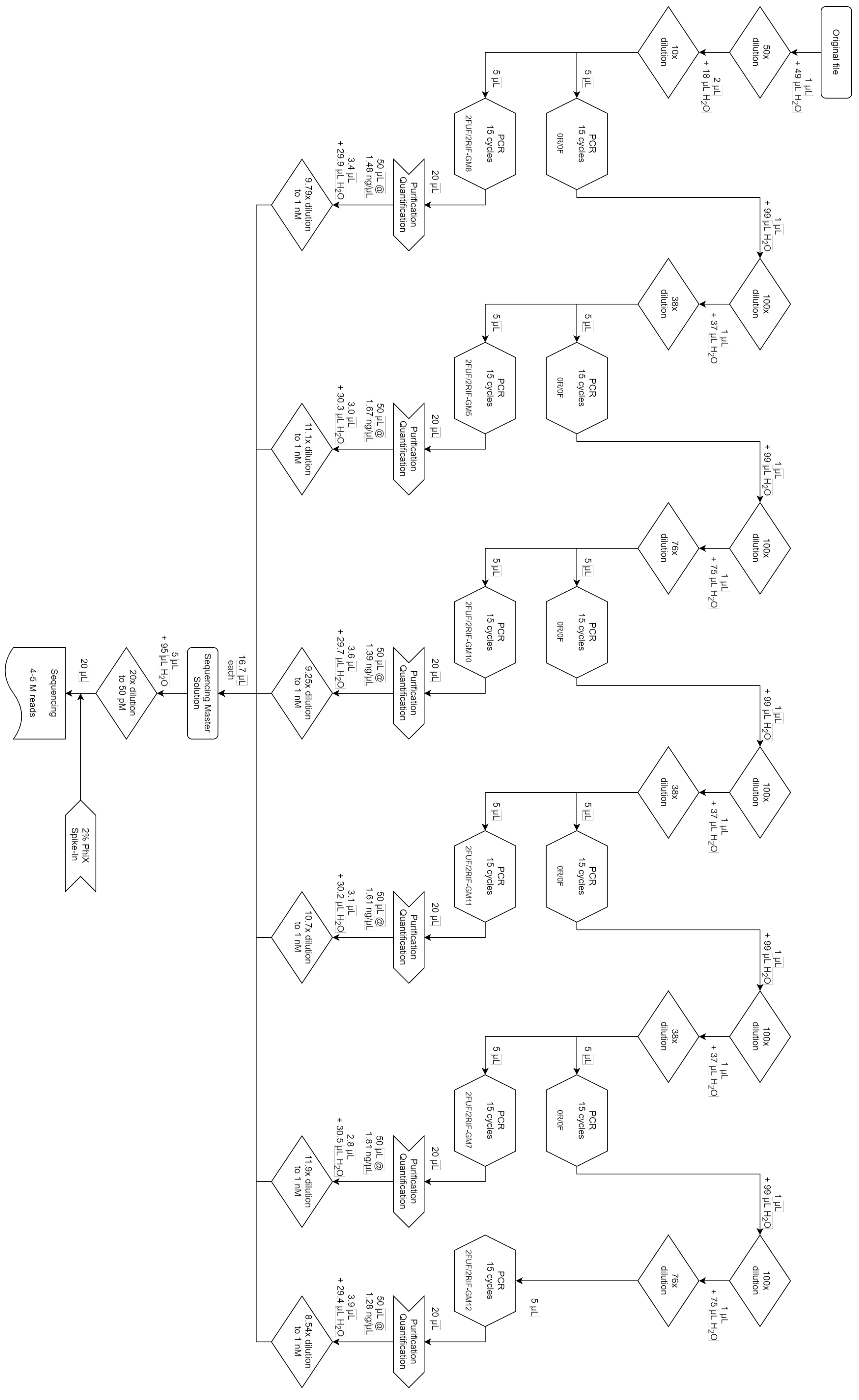


**Supplementary Figure 19: Experimental procedure for the amplification experiments using the GC-constrained oligonucleotide pool synthesized by material deposition.**

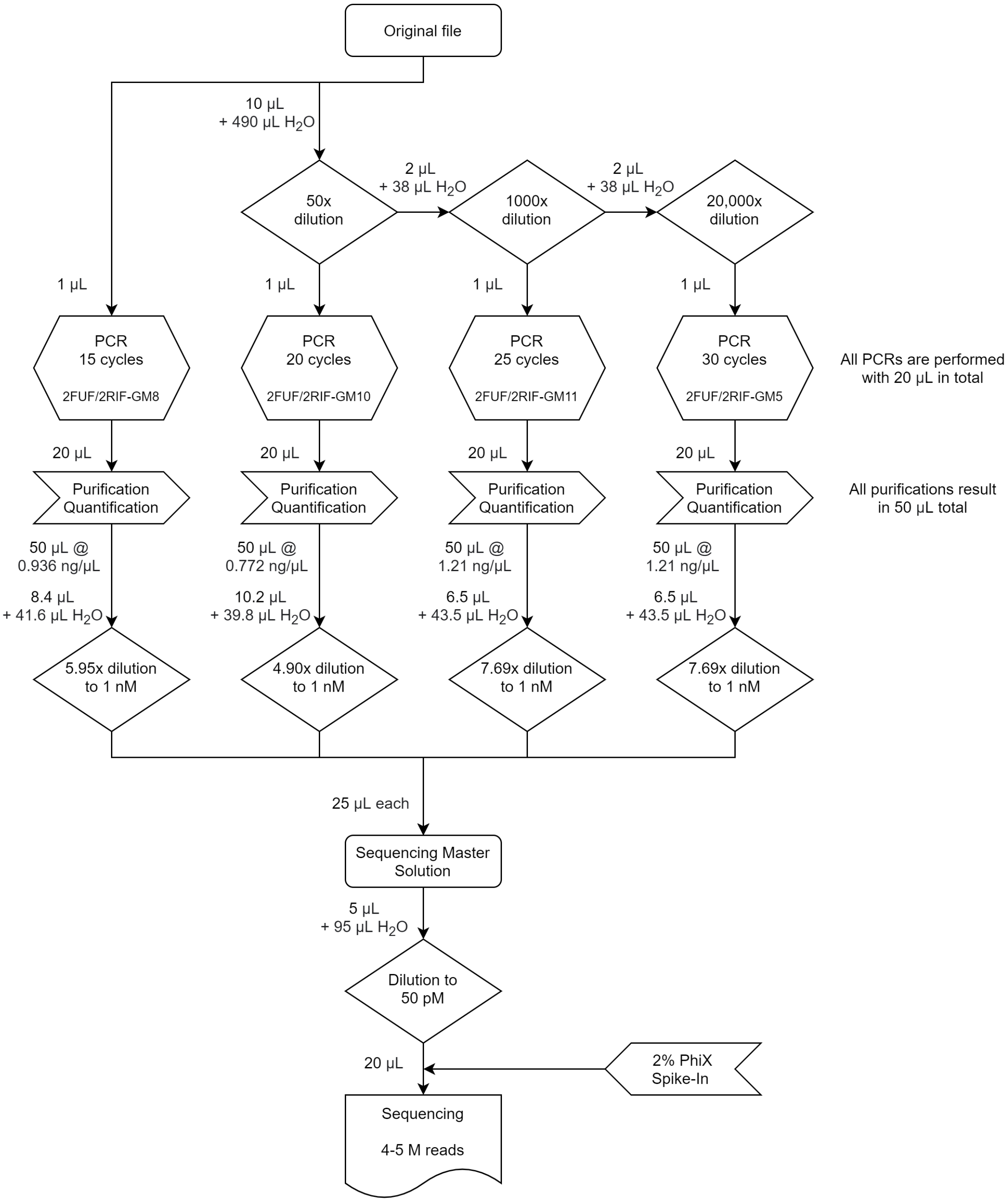


**Supplementary Figure 20: Experimental procedure for the amplification experiments using the unconstrained oligonucleotide pool synthesized by electrochemical synthesis.**

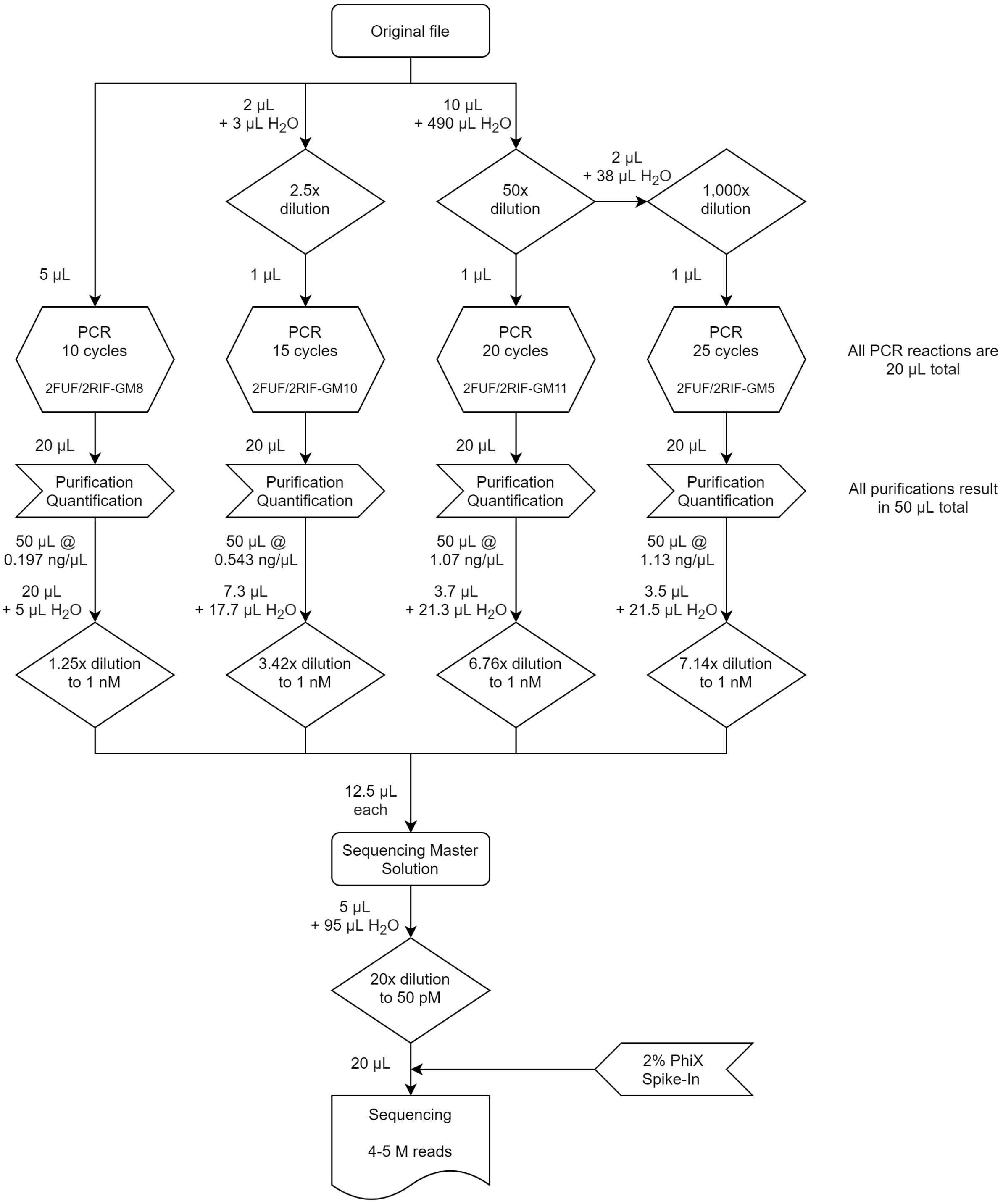


**Supplementary Figure 21: Experimental procedure for the amplification experiments using the GC-constrained oligonucleotide pool synthesized by electrochemical synthesis.**

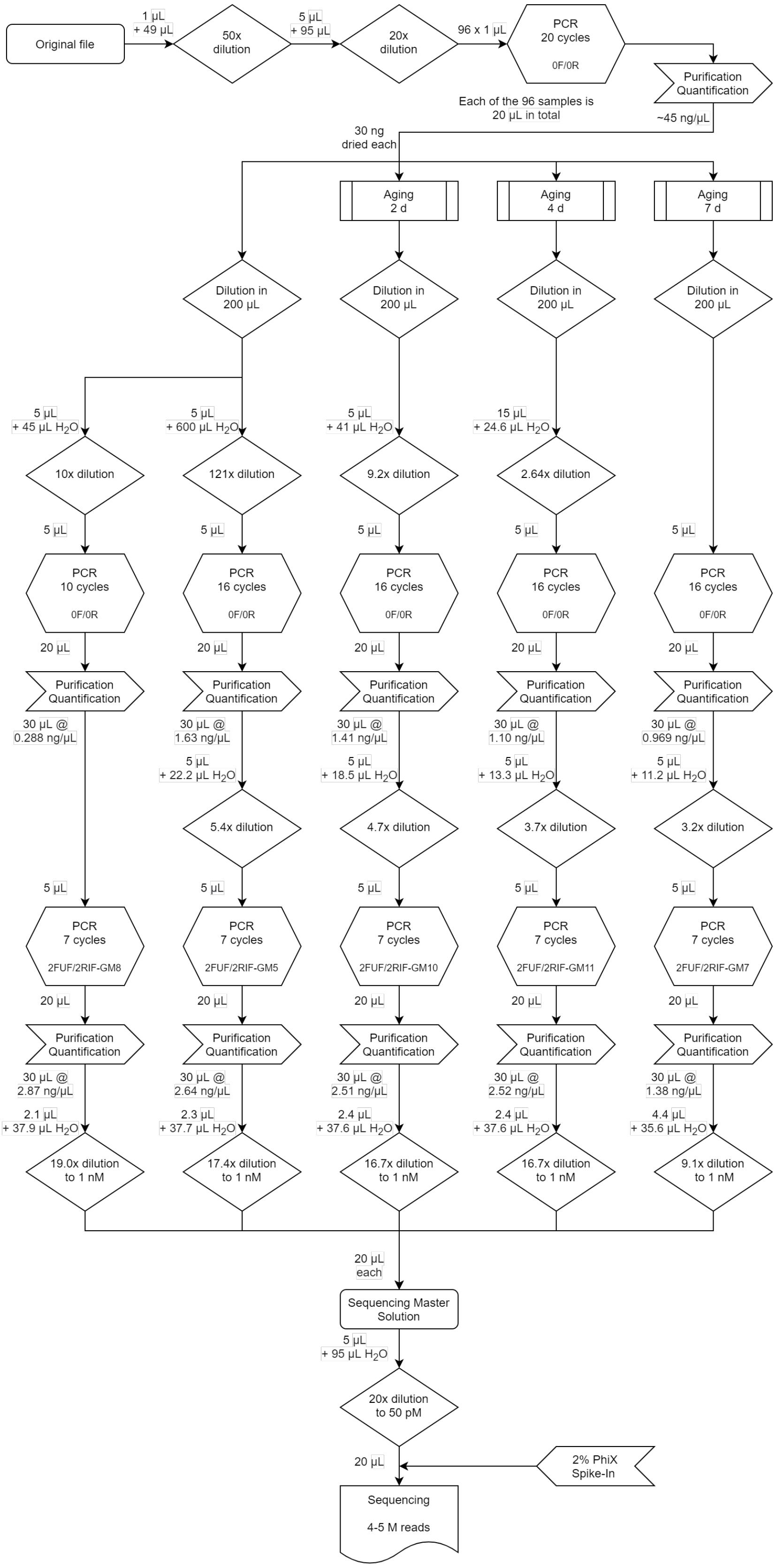


**Supplementary Figure 22: Experimental procedure for the storage experiments using the unconstrained oligonucleotide pool synthesized by material deposition.**

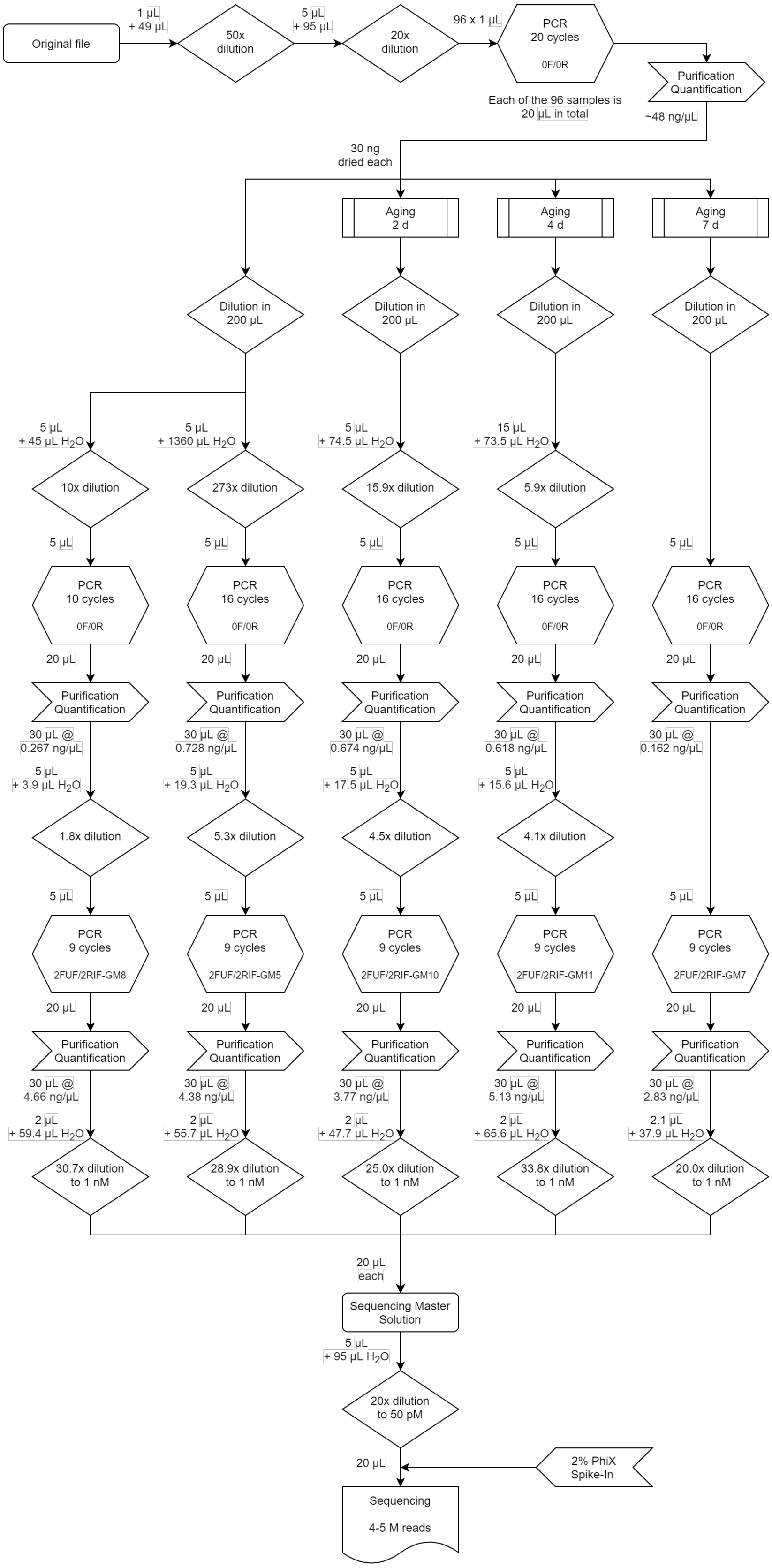


**Supplementary Figure 23: Experimental procedure for the storage experiments using the GC-constrained oligonucleotide pool synthesized by material deposition.**

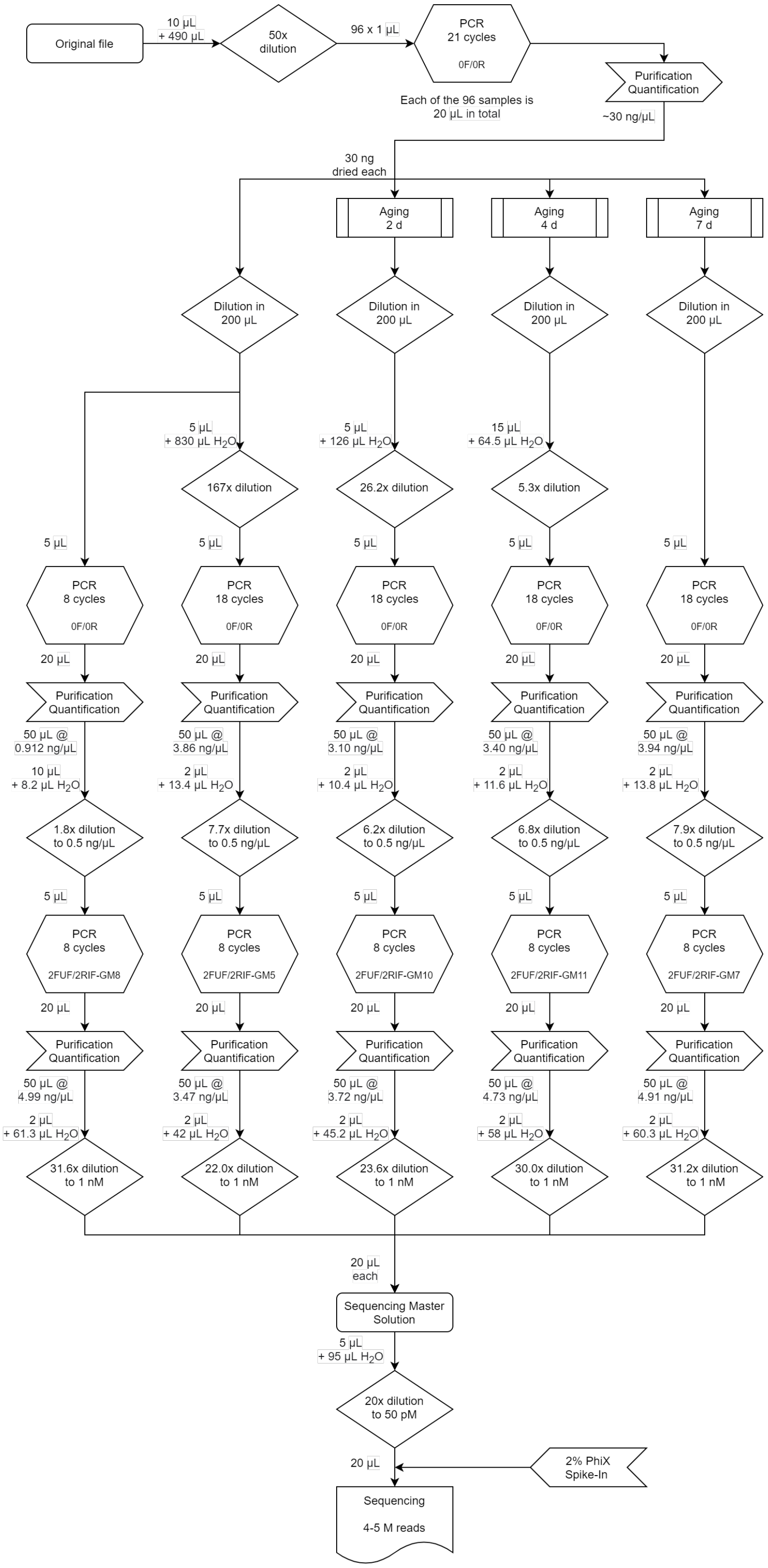


**Supplementary Figure 24: Experimental procedure for the storage experiments using the unconstrained oligonucleotide pool synthesized by electrochemical synthesis.**

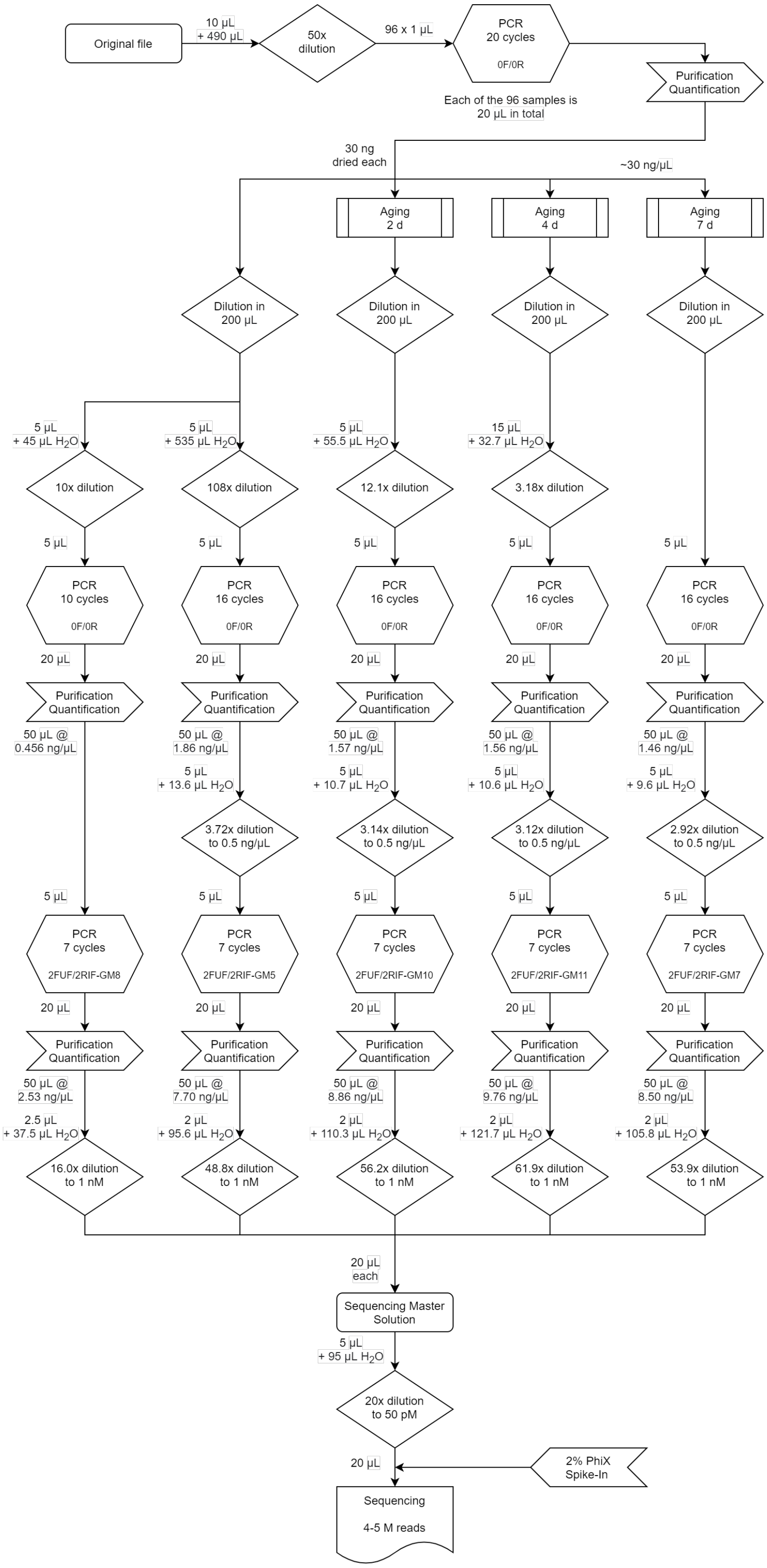


**Supplementary Figure 25: Experimental procedure for the storage experiments using the GC-constrained oligonucleotide pool synthesized by electrochemical synthesis.**

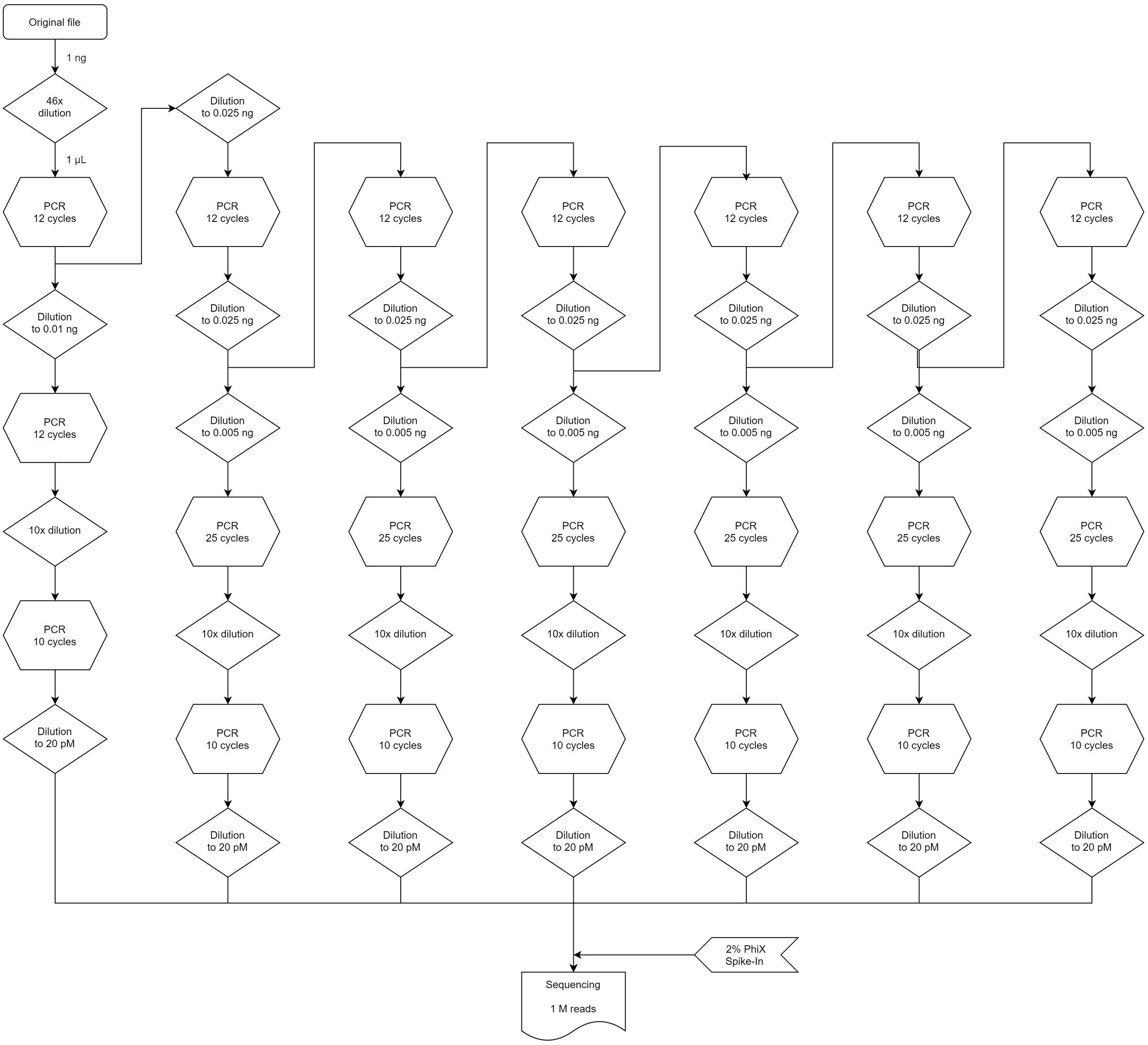


**Supplementary Figure 26: Experimental procedure estimated from the study of Koch et al.[4] for the generational experiments using a GC-constrained oligonucleotide pool synthesized by electrochemical synthesis.** For details, see Supplementary Note 5.
